# Self-Organization of Whole Gene Expression through Coordinated Chromatin Structural Transition: Validation of Self-Organized Critical Control of Genome Expression

**DOI:** 10.1101/852681

**Authors:** Giovanna Zimatore, Masa Tsuchiya, Midori Hashimoto, Andrzej Kasperski, Alessandro Giuliani

**Author notes:** Corresponding authors: Alessandro Giuliani (RQA and PCA); Masa Tsuchiya (SOC analysis).

## Abstract

Through our studies on whole genome regulation, we have demonstrated the existence of self-organized critical control (SOC) of whole gene expression - genomic self-organization mechanism through the emergence of a critical point (CP) at both the cell population and single cell level. In this paper, based on HRG and EGF-stimulated MCF-7 breast-cancer cell line, we shed light on the origin of critical transitions stemming from coordinated chromatin remodeling. In so doing, we validated the core of the SOC control mechanism through the application of a non-linear signal analysis technique (Recurrence Quantification Analysis: RQA), and of Principal Component Analysis (PCA). The main findings were:

1. Transcriptional co-regulation follows a strong and invariant exponential decay as between gene spacing along the chromosome is increased. This shows that the co-regulation occurs on a mainly positional basis reflecting local chromatin organization.
2. There are two main fluctuation modes on the top of the cell-kind specific gene expression values spanning the entire genome expression. These modes establish an autonomous genomic critical control system (genome-engine) through the activation of the CP for cell-fate guiding critical transitions revealed by SOC analysis.

The elucidation of the link between spatial position on chromosome and co-regulation together with the identification of specific locations on the genome devoted to the generalization of perturbation stimuli, give a molecular basis to the self-organization dynamics of genome expression and cell-fate decision.

## 1 Introduction

It is a fundamental challenge in current bioscience to elucidate the mechanism by which an integrated genome system guides cell fate change. This mechanism acts through the complex spatio-temporal self-organization of the genome involving on/off switching of thousands of functionally unique heterogeneous genes in a remarkably cooperative manner (MacArthur, 2009; Chang, 2006). However, there are two main fundamental physical difficulties in achieving such large-scale coordinated control on a gene-by-gene basis. The first one is the lack of a sufficient number of regulatory molecules in a cell to reach a stable thermodynamic state (i.e., breakdown of the mass action law). The low copy number of specific gene mRNAs provokes stochastic noise (Wolynes, 2010), thereby inducing a substantial instability of genetic product concentrations falsifying any gene-by-gene feedback control hypothesis (Raser, 2005; Yoshikawa, 2002).

The second one derives from the huge linear dimension of human DNA molecule (around 2 meters) with respect to cell nucleus that makes chromatin very far from a Turing-like string freely accessible by regulatory molecules at single gene level.

This extreme compression generates by a complex folding of chromatin generating a fractal-like organized structure at different orders of magnitude (Cremer, 2004). There are many evidences of the existence of cell type specific chromatin folding organization (Mayer, 2005; Zentgraf, 1994) as well as of transcription dependent local changes in chromatin organization (Bohn, 2010). In this work, we explore the consequences on transcription dynamics of considering chromatin remodeling as the main material basis of gene expression regulation.

Our recent studies (Tsuchiya, 2015, 2016, 2019; Giuliani, 2018) demonstrated Self Organized Criticality (SOC: see Supplementary file A for basic SOC principles and Supplementary file B for essential summaries of SOC results in MCF-7 cell responses) is a physically motivated candidate for such a goal. The basic issues of genome expression control by SOC are:

1. Whole genome expression is dynamically self-organized through the emergence of a critical point (CP) at both the cell-population and single-cell levels - self-organized critical control (SOC) of whole expression,
2. The CP represents a specific set of critical genes, which has an activated (ON) or deactivated (OFF) state. The ON-OFF switch of the CP state occurs through change in critical transition on its singular behaviors. Furthermore, this set of critical genes acts as genome-attractor. This implies that any random sampling (more than 50 genes) of stochastic genome expression converges to the dynamics of the CP,
3. In OFF state of the CP, the stochastic perturbations propagate locally but when the particularity of the disturbance activates the CP, the perturbation can spread over the entire system in a highly cooperative manner,
4. As the system approaches its critical point, CP is ON, and global behavior emerges in a self-organized manner. In such a way is possible that an autonomous critical-control genomic system (*genome-engine*) develops through the formation of dominant cyclic flux between critical states. Coherent perturbation on the genome engine through the activation of the CP drives cell-fate decision.

While such a model is able to predict many observed features of biological regulation at both single cell and population levels (Giuliani, 2019), it still lacks a foundation in terms of molecular mechanisms involved. In this paper, we give a proof both of the local character of the fine-tuning regulation along the chromosome and evidence of SOC critical states at single gene expression level. Both issues point to chromatin flexibility around its cell-kind specific organization as the main driver of gene expression regulation at large. We faced the above tasks by a non-linear signal analysis technique (Recurrence Quantification Analysis, see Supplementary file C) and by a classical statistical technique (Principal Component Analysis) as applied to time series of transcriptome data in two paradigmatic conditions.

The present work deals with this proof and redounds around three main issues (as the three paragraphs in results):

i. To give a proof-of-concept of the existence of a relation between spatial proximity of genes along the chromosome and their co-regulation (see par. 3.1).
ii. To investigate the presence of specific genes (‘hot-spots’) organizing the fluctuations around the ‘cell-kind specific ideal expression value’ (‘attractor’) and driving SOC mechanism by means of self-organization (see par. 3.2).
iii. The clarification of positional vs. functional character of gene expression regulation (see par. 3.3).

## 2 Materials and Methods

### 2.1 The biological case

The biological case studied in this work comes from (Saeki, 2009) and has optimal characteristics to characterize the essential features of biological regulation. Actually, it allowed us to derive the essential of SOC based gene expression regulation reported in previous papers (Tsuchiya, 2014, 2015)

The data set collects gene expression data relative to the same cell-kind (MCF7 a breast tumor immortalized line) observed for 18 time points after the administration of two different stimuli: Epidermal Growth Factor (EGF) and Heregulin (HRG). In EGF-stimulated cells, cancer cell develops cell proliferation without cell differentiation, whereas HRG-stimulation induces cell differentiation (Nagashima, 2007). Notably, while EGF does not provoke a critical transition in MCF7 cells, HRG does (Tsuchiya, 2019).This fact allows a direct comparison between the two EGF and HRG conditions to single out what discriminates the two situations in presence of a fully controlled and homogenous system (same cell-kind, time points, and microarray system).

### 2.2 General considerations and data analysis strategy

The common experience of any scientist working with transcriptome microarrays is the near to unity correlation between gene expression profiles of independent samples coming from the same cell-kind (see Fig. 1 in Tsuchiya, 2016)

**Figure 1.**
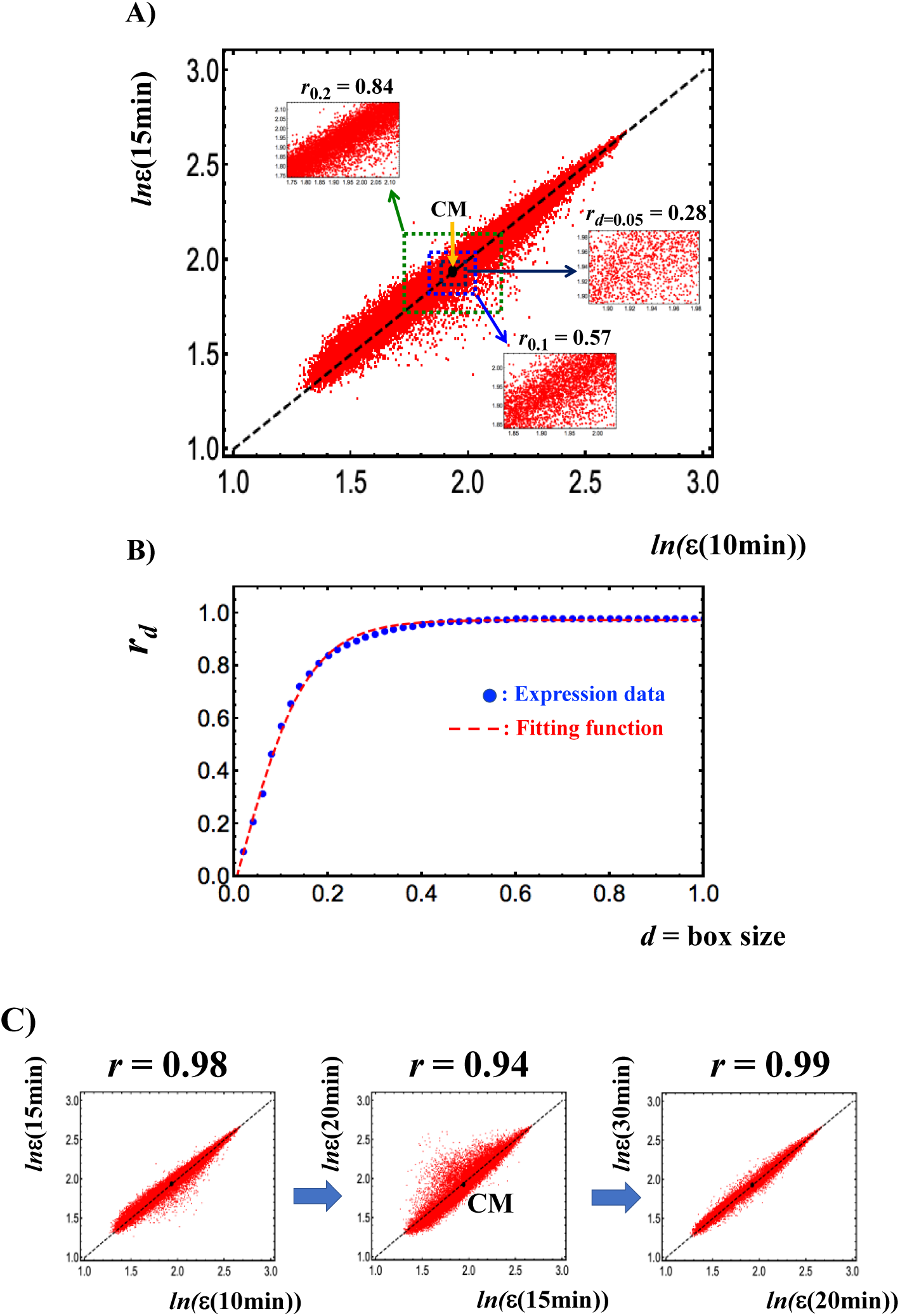
Transition from a stochastic to a genome-wide attractor profile. **A)** The figure reports the Pearson’s correlation between two independent samples **(10min vs. 15min)** of MCF7 (axes of the plot) in terms of expression levels of around 23000 mRNAs (vector points). The overall correlation is near to unity (*r* = 0.98) consistently with the existence of a main attractor correspondent to the cell-kind. At smaller scale of variation, the correlation (*r*_*d*_) decreases, for the increase in relative importance of gene expression fluctuations. **B)** The attractor structure is evident starting from a variation range (box size) *d* = 0.45. From that scale onward, between expression profiles correlation *r*_*d*_ follows a tangent hyperbolic function as signature of transition from a stochastic to a genome-wide attractor profile. **C)** The correlation between gene expressions profiles at different time points reveals the occurrence of critical transition at 15-20 min around the CM (back solid dot) correspondent to a decrease in Pearson r and visually to an increase scattering of points from the identity line. A tipping point in terms of the critical transition exists at 15min (see more in Figure 5). This stems from the (coil-globule) critical transition at the CP, where the CP corresponds to the CM when expression is ordered by *nrmsf*.

Figure 1A refers to MCF-7 cell line and clarifies this point: the axes refer to two independent MCF7 samples whose single gene expression values are the points of the graph (around 23000, expression values in logarithm units), the *d*-value corresponds to the range (box size) of variation, inside which the correlation (Pearson coefficient, *r*) is computed. The correlation computed overall is near to unity (*r* = 0.98) and declines at decreasing range of variation. Figure 1B shows the reaching of a plateau correlation at *d* = 0.45. This remark outlines how correlation values are tightly dependent on the observation scale. The scale dependence of the correlation is instrumental to keep alive both the functionality of the tissue (the specialized physiological function asks for an invariant “ideal” pattern of gene expression) and the flexibility required to adapt to changing microenvironment, tuning the specific gene expressions at the small scale. This fine-tuning does not alter the global profile invariance and corresponds to the scattering of the points around the identity line (*y = x* line in Figure 1A). The dispersion across the identity line corresponds to the equilibrium around a definite physiological (tissue dependent) state that, as expected in SOC corresponds to a ‘critical dynamically stable state’ encompassing continuous fluctuations around the cell-kind specific profile.

This general property of gene expression regulation tells us that the main driver of gene expression is the ‘cell-kind’. Humans have approximately 400 cell-kinds (Vickaryous, 2006), each one correspondent to a ‘global stable expression pattern’ out of the infinite number of patterns possibly stemming from around 25000 genes each potentially varying across four order of magnitude. This is the image in light of the limited number of allowed ‘chromatin global organization patterns’ each corresponding to a cell-kind attractor of transcription dynamics.

#### 2.2.1 Pre-processing data: normalization

In our formalization, the expression data matrix has genes (22035) as statistical units (rows) and the expression profiles at different times (18) as variables (columns).

On a statistical point of view, any effort for demonstrating the existence of a relation between spatial proximity of genes along the chromosome and their co-regulation must eliminate the ‘batch effect’ we referred in the previous paragraph stemming from a specific cell-kind that masks the fluctuations within cell-kind attractor.

This makes mandatory to approach the problem by preliminary normalization of the single gene expression. This row-by-row (genes are the rows of the original matrix, time points are the columns) normalization was faced transforming expression values to z-scores by the formula:

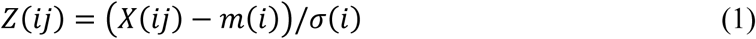

where *X*(*ij*)-is the expression value of gene *i*, at time *j* (*t*j), and *m*(*i*)-and *σ*(*i*)-the average and standard deviation respectively of *X*(*i*)-across all the times.

After this transformation, we posit two A and B genes are co-regulated (at a specific time *j*) if they have very similar z-scores. In other words, two A and B genes are co-regulated if they have approximately the same difference with respect to their respective cell-kind ideal expression value. The order of genes along the chromosome is mathematically equivalent to a discrete time-series whose values are the single gene (normalized) expression values sampled on chromosome location (gene, gene2, gene3 according to the relative order along the chromosome). This allows approaching the problem of the link between gene co-regulation and spatial position on chromosome by means of signal analysis methods. The demonstration of a link between the spacing of genes along the chromosome and the probability to be co-regulated is thus a proof-of-concept of the supposed equivalence between chromatin local flexibility and gene expression variability (i.e., temporal expression variance: *nrmsf*) at the basis of our previous works (Tsuchiya 2016, 2019).

#### 2.2.2 Principal Component Analysis (PCA)

The presence of a cell-kind specific gene expression profile (Figure1) has very important consequences for the Principal Component Analysis (PCA) based approach to regulation. If the system stays in the very center of the attractor, with no fluctuation, we observe a maximal correlation between the variables because each time profile is identical to all the others. This implies all the variance of the system lays along the ideal line ordering the genes from the less to the more expressed. We call this ideal line *‘cell-kind attractor’*, this line corresponds to the first principal component (PC1). Consistently to what depicted in Figure 1, PC1 (see Results) explains more than 90% of total variance.

We can factorize total variance (how diverse the values in the matrix are) as ‘Between Genes Variance’ and ‘Within Gene Variance’:

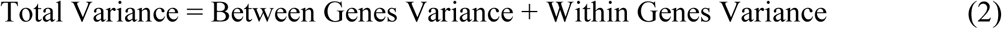

Normalized root-mean-square fluctuation (*nrmsf*), the parameter we used in previous works (Tsuchiya 2016, 2019) to group genes in different critical states, has to do with the second term of the equation (2): it comes from es the variance internal to each gene (i.e. each gene has a specific *nrmsf* value). In other terms, *between* variance is relative to variability along the columns (i.e., variance of whole gene expression at a specific time), *within* variance is relative to variability along the rows (i.e., expression variance of a gene over the time points). When in presence of a dominant first component (size component, Jolicoeur, 1960), the fluctuations around the attractor are accounted for minor components (from second onward) that, by construction are orthogonal to each other and to the main order parameter (PC1).

Recurrence Quantification Analysis (RQA), the signal analysis procedure (Marwan, 2007) we adopted to deal with the gene co-regulation along the chromosomes, deals with this minor fluctuation on the top of cell-kind attractor.

Near a tipping point (see Figure 1, panel C), the second term of (2) becomes increasingly important: some genes deviate ‘more-than-usual’ from their ideal expression level in direction(s) orthogonal to PC1 (we discovered two main directions of this motion: PC2 and PC3, see Results). This provokes a decrease in between profiles correlation especially evident in ‘high motion’ (super critical in SOC terminology) genes (profile correlation going from 0.98 to 0.75 for supercritical state, Fig.4 in (Tsuchiya, 2015).

**Figure 2.**
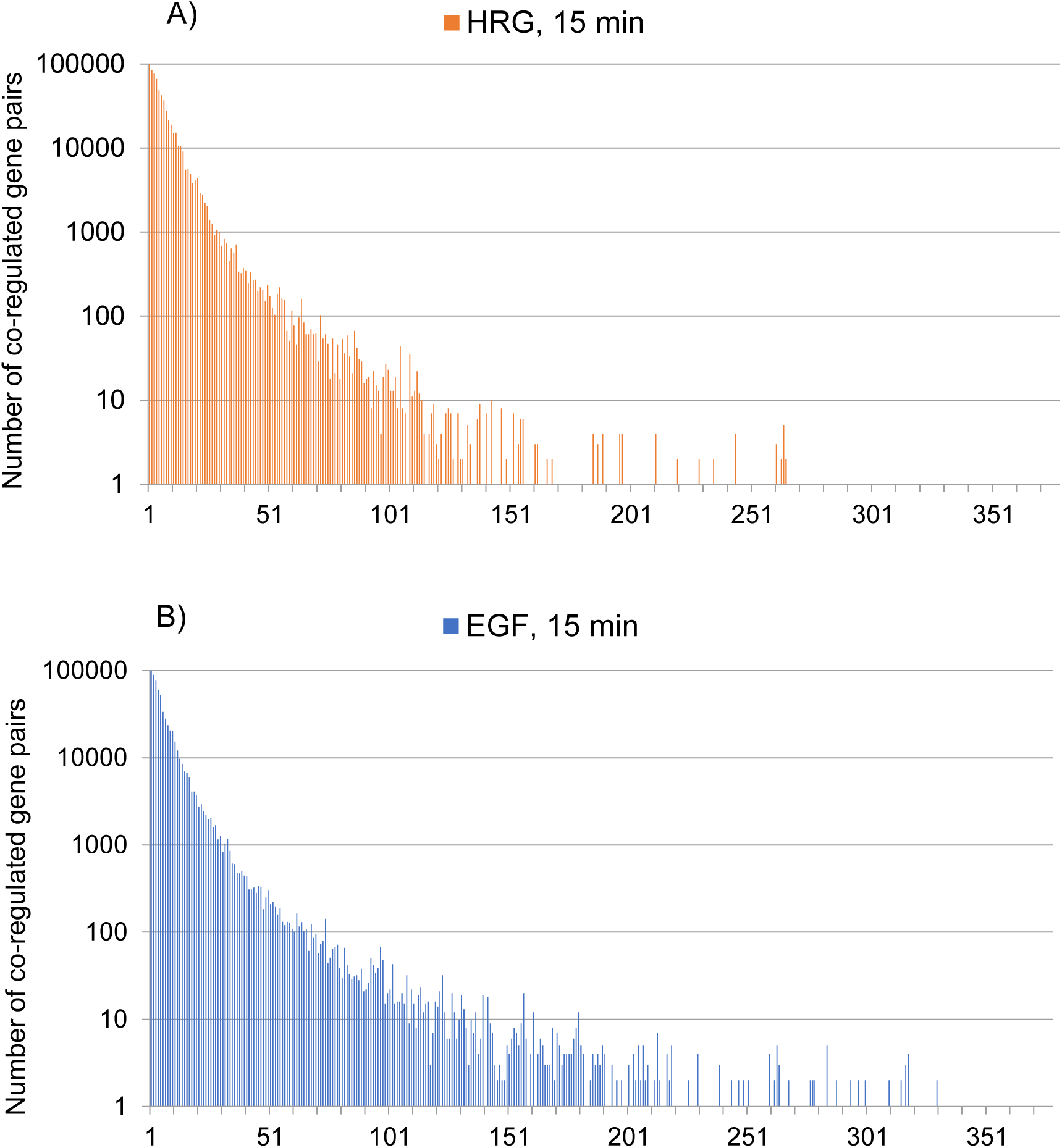
Recurrence spectrum along the chromosome 1: The graphs are relative to HRG (top) and EGF (bottom) conditions as for time 15 min and chromosome 1. The y-axis corresponds to the number of recurrences (number of co-regulated gene pairs). The x-axis corresponds to the spacing of recurrent couples along the chromosome. The spacing is expressed in terms of number of genes between the recurrent pairs. The exponential decay of co-regulation entity with increasing between genes spacing is identical for different times of observation, different chromosomes and for the two EGF and HRG conditions. The local character of fine co-regulation around cell-kind specific value is the same for all the experiments.

**Figure 3:**
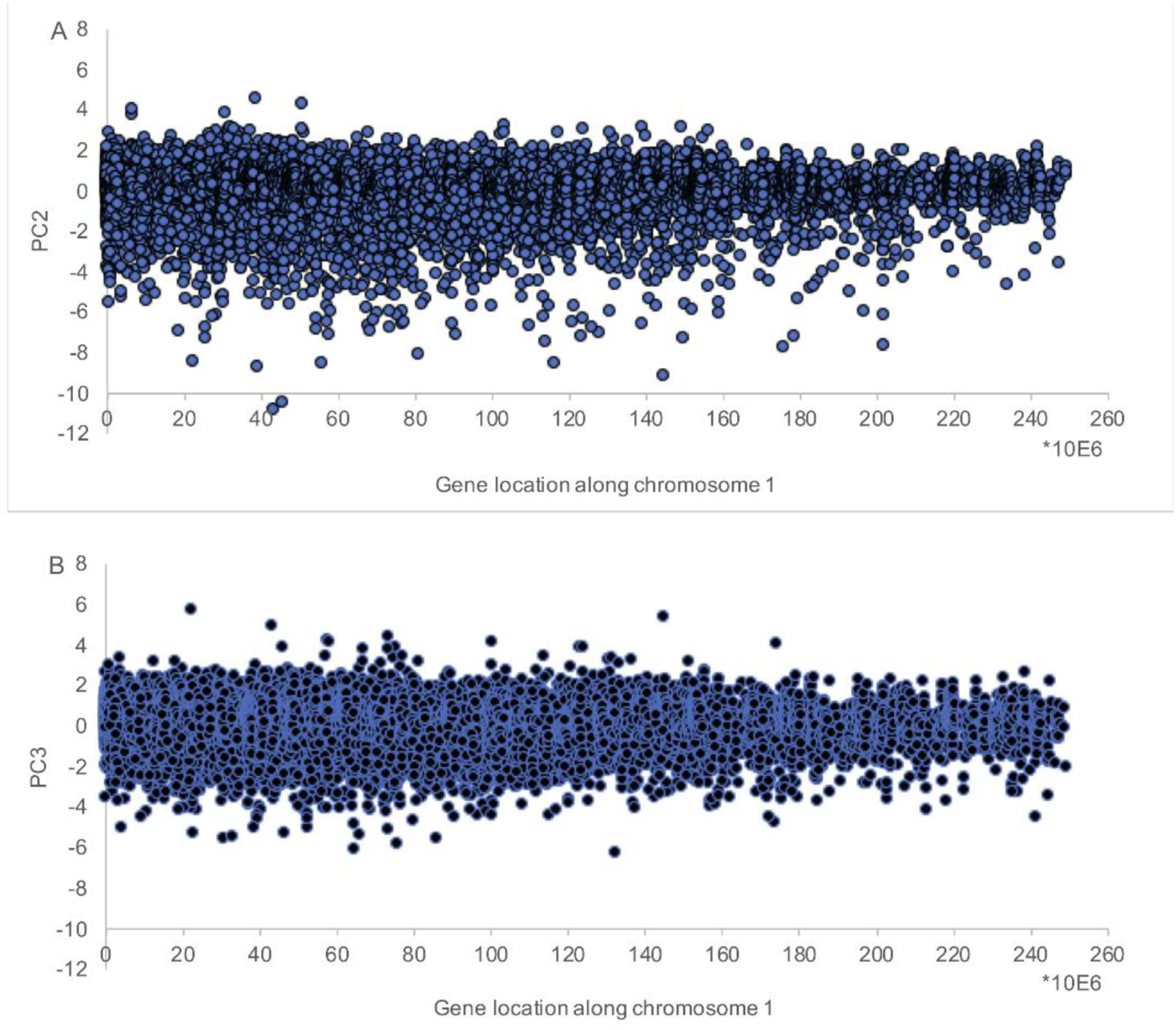
Distribution of PC2 and PC3 scores along chromosome 1. The x-axis is the gene location along chromosome 1, while the y-axis reports the correspondent PC2 (top panel) and PC3 (bottom panel) scores. The figure refers to chromosome 1 EGF at 15 minutes, but the same general pattern is identical across all the analyzed conditions.

**Figure 4:**
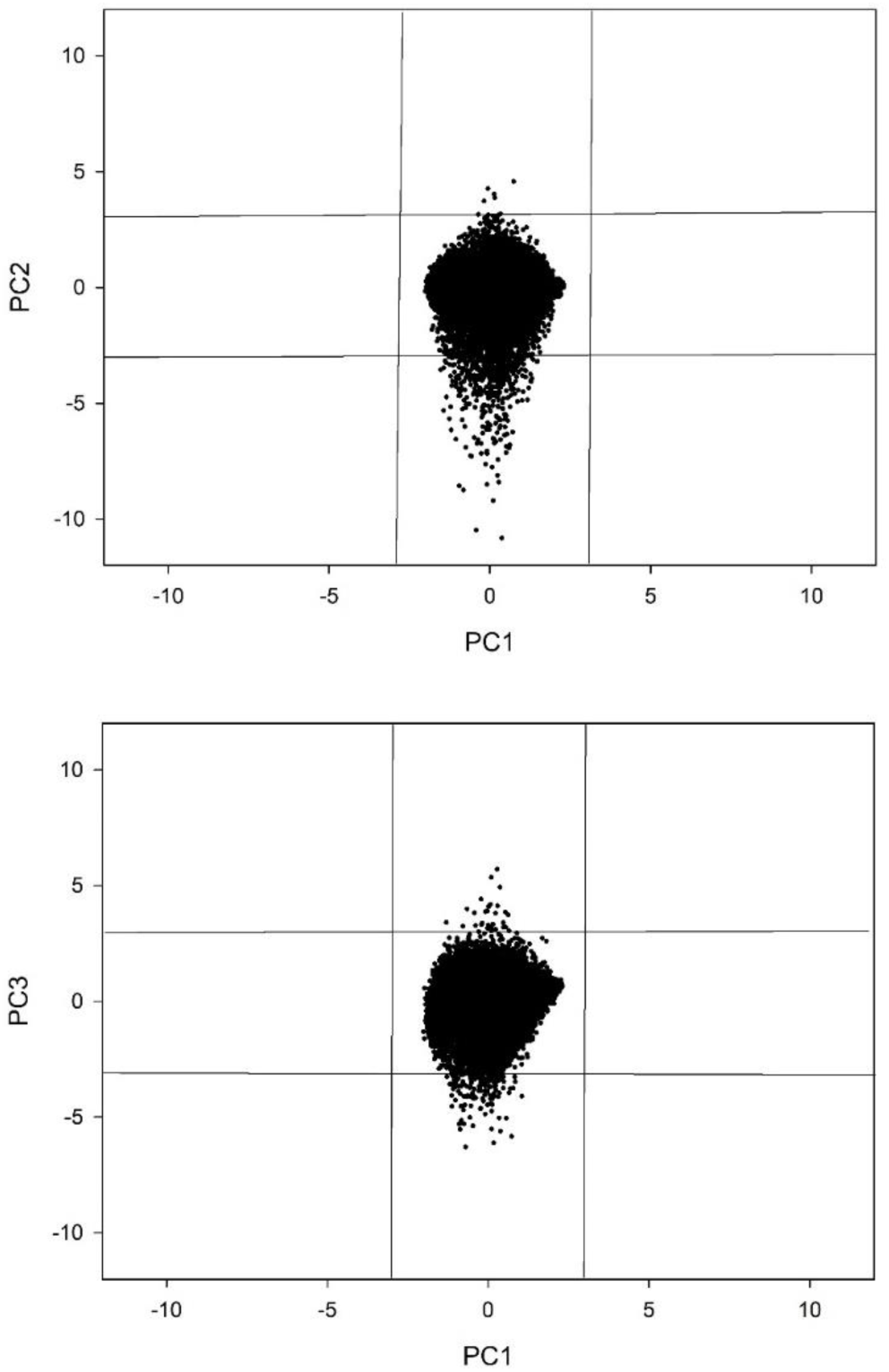
Distributional properties of fluctuations around cell-kind specific expression. The figure reports the projection of the genes in the PC1-PC2 (top) and PC1-PC3 (bottom) spaces. Vertical and horizontal lines correspond to the 3 Standard Deviation interval of PC1 and (PC2, PC3) respectively. Genes outside PC2 (or PC3) confidence intervals are the ‘hot spots’. All the components have by definition zero mean and unit standard deviation but while the variance of fluctuation modes (PC2, PC3) is driven by the outliers, the cell-kind attractor mode (PC1) has a continuous Gaussian distribution restricted into a much narrower variation range (no outliers are observed).

In PCA terms, the expression value *E*(i), for each gene *i*, across the 18 time points, can be reconstructed as a linear combination of the component scores as:

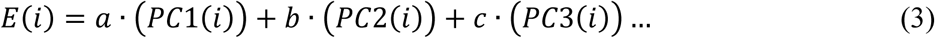

The equation (3) is intended on the entire set of 18 components but, if we stop the weighted sum at the first three components, thanks to the very strong correlation structure of the system, we can reconstruct the actual expression value with 99% of accuracy (see Results). PC2 and PC3 are coordinated ‘waves of fluctuation’ spanning the entire genome and we will see in the following that they account for the collective gene expression oscillations responsible for SOC. In the same heading, we expect that, in the case of a real transition (HRG) the relative importance of fluctuation (PC2 and PC3) modes should be greater with respect to the no transition (EGF) case.

#### 2.2.3 Functional classification by PANTHER

We checked the existence of a specific ‘functional signature’ of genes participating to cell-fate transition by PANTHER Gene Ontology software: (http://pantherdb.org/). This software takes as input a list of genes and produces as output the distribution of the genes in different functional classes. This happens by means of two different ontologies correspondent to ‘molecular function’ and ‘biological processes’ to which the genes are associated. In the case of ‘molecular function’, the genes are partitioned across 9 classes: *transporter, translation regulator, cargo receptor, transcription regulator, catalytic, molecular function regulator, molecular transducer, structural, binding*.

The ‘biological process’ spectrum has 14 categories: *biogenesis, cellular process, localization, reproduction, biological regulation, response stimulus, development, multi cellular organismal, adhesion, metabolism, cell proliferation, immune system, biological phase, rhythmic*. Both ‘molecular function’ and ‘biological process’ ontologies are not exclusive: the same gene can be part of more than one category. The 200 elements gene lists correspondent to different dynamical roles were described in terms of their PANTHER spectra to identify some eventual enrichment in a specific function or process.

## 3 Results

### 3.1 Spatial proximity of genes and co-regulation

The mono-dimensional series correspondent to normalized expression values of the genes ordered along their relative position on relative chromosomes, was analyzed by means of Recurrence Quantification Analysis (RQA), a widely used non-linear signal analysis technique (see Supplementary file C for a brief explanation of RQA) particularly fit for non-stationary series (Webber&Zbilut, 1994; Giuliani, 1998) and successfully used in a multitude of disciplines from physiology (Zimatore et al, 2011, 2015) to earth science (Zimatore et al, 2017) and finance (Orlando&Zimatore, 2017; Zimatore et al, 2018). In this work we relied on percent of recurrence (REC) descriptor that was expressed in terms of the spectrum of recurrence distribution at different separation along the sequence (RQI): the y corresponds to the number of recurrences scored between genes located at a distance x (expressed as number of genes between i and j recurrent points) (see Figure 2).

The recurrence spectra (each recurrent event correspond to a pair of co-regulated genes) depict a very robust exponential decay of co-regulation with increased spacing of gene pairs along the chromosome. This behavior is consistent with the effect of local structural chromatin features (folding status, flexibility) on gene co-regulation. Figure 2 reports the recurrence spectrum (radius = 60% of mean difference, see Supplementary file C) along chromosome 1 at T1 (0 min) and T3 (15 min) for HRG and EGF conditions. The same pattern holds for all the time points, conditions and chromosomes.

This demonstrates that transition and no-transition conditions do not imply qualitatively different regulation mechanisms: the response to the two stimuli happens through the same basic mechanism involving local chromatin folding status, the differences are only quantitative (and very subtle) as we will discover in the following.

The local character of co-regulation is consistent with the ‘domino-effect’ of SOC sandpile model, where each sliding ‘grain of sand’ has a direct effect on the neighbor and the formation of ‘big avalanches’ depend on the ‘traveling’ of local perturbation by chaining ‘single step’ events (see Supplementary file B for a brief explanation of SOC). The occurrence of genome-wide global perturbation under the exponential decay of co-regulation suggests that the chaining ‘single step’ follows first order transitional behavior on chromatin Yoshikawa, 1996, 2002; Zinchenko, 2008). The dependence of co-regulation on the location on chromosome gives a proof-of-concept of main hypothesis at the basis of our previous works: gene expression fluctuations mirror chromatin flexibility. This is consistent with the observation that chromatin modifications can cause a cascade of events leading to promote or block gene expression (van der Knaap, 2016).

This fact in turn sets the fundamental premises for a mainly ‘position’ as opposed to a mainly ‘function’ based regulation, as we will discuss in the following.

The last point of this section is a sort of ‘disclaimer’ about the (largely unescapable) terminological confusion of a research work located at very edge between Physics and Biology. Each specific discipline attaches to the same term a slightly different meaning, the *de novo* introduction of ‘new terms’ for research on the ‘edge’ is clearly unfeasible, so we prefer to overly declare the possible ambiguities linked to the use of crucial terms frequently encountered in the paper:

1. **Attractor**: We used this word in both a *general* ‘a preferred configuration the system tends to’ and a more *specific* ‘a stable (albeit critical) state actively maintained by system fluctuation’ sense. The first case is more familiar to the biologist: it simply describes the evidence that each cell-kind has a typical profile of gene expression spanning the entire genome. The most straightforward biological analogy is the existence of a protein ‘native structure’ that remains recognizable along the (small) fluctuations due to thermal motion (molecular dynamics). In this sense, all the different genes participate of the same attractor whose strength is evident by the fact that the by-far major part of system variance is accounted by PC1. The second, more physically oriented, meaning of the word ‘Attractor’ has to do with SOC model. In this case, the term refers to the contemporary presence of three (‘super-critical’, ‘near-critical’ and ‘sub-critical’) expression states that tend to the ‘critical state’ (attractor). In this frame, the attractor has a main ‘core’ correspondent to the center of mass (see Supplementary file B) genes and a periphery correspondent to the most fluctuating (in the following ‘hot-spots’) genes. These two different notions are mutually consistent; in the following, we will demonstrate the center of mass genes are those with an overwhelming contribution of PC1 (near zero value for PC2 and PC3 ‘fluctuating’ components).
2. **Ideal**: The attribute ‘Ideal’ has no semantic implication, it is only used (like frequently happens in biomedical literature) to identify a ‘paradigmatic’ or ‘exemplar’ configuration. In this case, the ‘ideal cell-kind profile’ corresponds to what we called ‘attractor’ in the generic sense. Each gene has an ‘Ideal’ (cell-kind specific) expression value that in our case corresponds to its mean value across the 18 time points.
3. **Motion:** PCA corresponds to the ‘eigenvalue/eigenvector’ decomposition of classical dynamics. This means that the displacement from an average value can be equated to the ‘motion’ of a material point. The variance of a probability distribution is analogous to the moment of inertia (classical mechanics term) of a corresponding mass distribution along a line, with respect to rotation about its center of mass. The fact PCA solution maximizes the ‘explained variance’ of a multivariate data set by the projection of the original data set into a new space spanned by a minimal number of independent axes corresponds to say PCA solution maximizes the moment of inertia. Here we use ‘variance explained’ and ‘entity of motion’ in an interchangeable way.

### 3.2 Hot spots, critical point and transition

Table 1 reports the results of PCA as applied to EGF condition in terms of both eigenvalue distribution (Table 1a) and component loadings (correlation coefficient between original variables and components, Table 1b).

**Table 1a.** Eigenvalues distribution (EGF)

| Component | Eigenvalue | Explained Variance (%) | Cumulative variance (%) |
| --- | --- | --- | --- |
| *PC1* | **17.767** | **98.71** | **98.71** |
| *PC2* | **0.073** | **0.41** | **99.12** |
| *PC3* | **0.046** | **0.26** | **99.38** |
| *PC4* | 0.027 | 0.15 | 99.51 |
| *PC5* | 0.017 | 0.10 | 99.60 |

**Table 1b.**
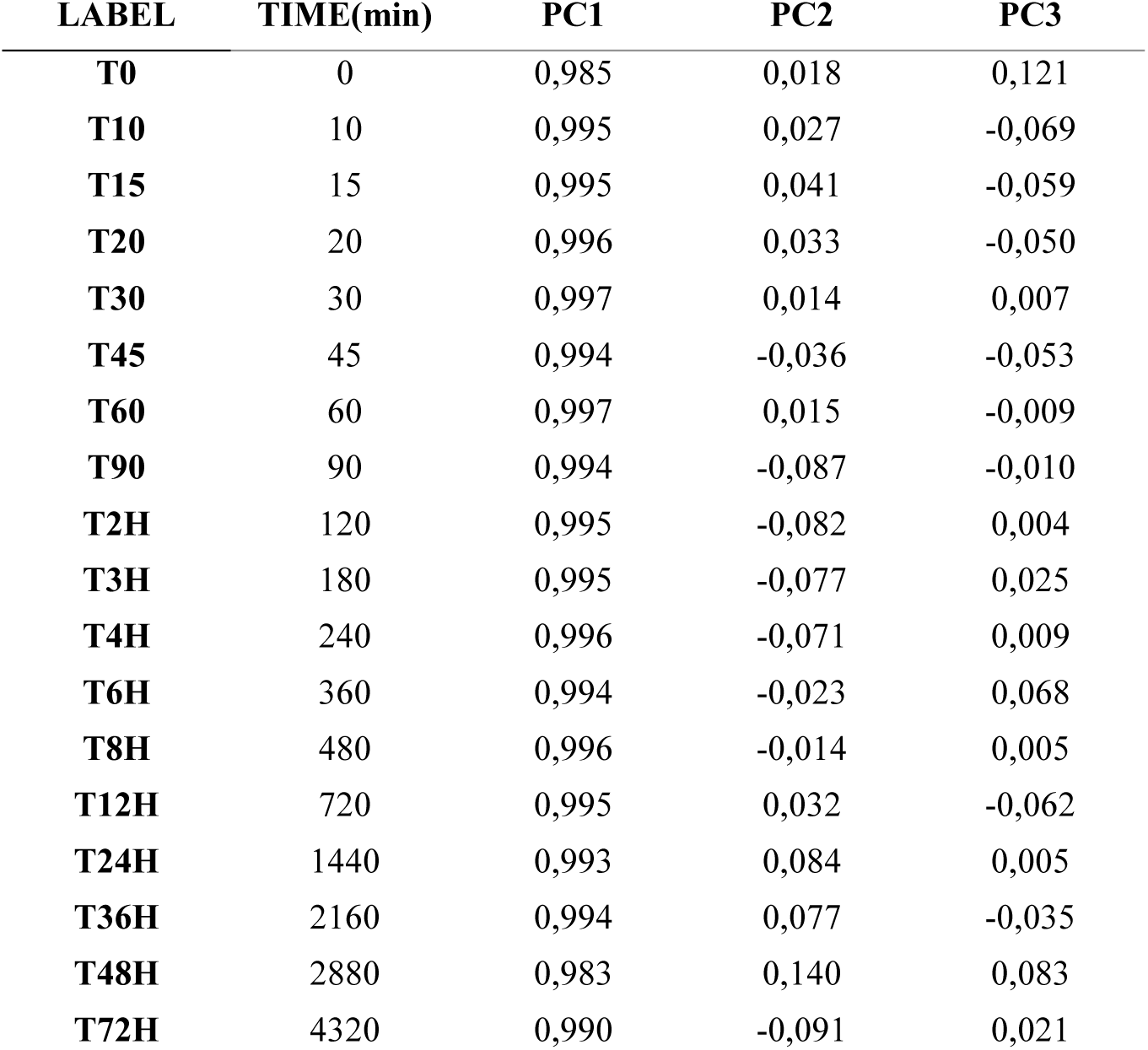
Loading Pattern (EGF)

| LABEL | TIME(min) | PC1 | PC2 | PC3 |
| --- | --- | --- | --- | --- |
| **T0** | 0 | 0,985 | 0,018 | 0,121 |
| **T10** | 10 | 0,995 | 0,027 | -0,069 |
| **T15** | 15 | 0,995 | 0,041 | -0,059 |
| **T20** | 20 | 0,996 | 0,033 | -0,050 |
| **T30** | 30 | 0,997 | 0,014 | 0,007 |
| **T45** | 45 | 0,994 | -0,036 | -0,053 |
| **T60** | 60 | 0,997 | 0,015 | -0,009 |
| **T90** | 90 | 0,994 | -0,087 | -0,010 |
| **T2H** | 120 | 0,995 | -0,082 | 0,004 |
| **T3H** | 180 | 0,995 | -0,077 | 0,025 |
| **T4H** | 240 | 0,996 | -0,071 | 0,009 |
| **T6H** | 360 | 0,994 | -0,023 | 0,068 |
| **T8H** | 480 | 0,996 | -0,014 | 0,005 |
| **T12H** | 720 | 0,995 | 0,032 | -0,062 |
| **T24H** | 1440 | 0,993 | 0,084 | 0,005 |
| **T36H** | 2160 | 0,994 | 0,077 | -0,035 |
| **T48H** | 2880 | 0,983 | 0,140 | 0,083 |
| **T72H** | 4320 | 0,990 | -0,091 | 0,021 |

It is worth noting the ‘cell-kind attractor’ accounts for around 99% of total variance (PC1), while only two minor components (PC2, PC3) can be considered as *bona fide* signals. Figure 3 reports the distribution of PC2 and PC3 scores along the chromosome 1.

From Figure 3, it is evident the presence of ‘outliers’, genes whose expression is driven by the oscillatory PC2 and PC3 modes much more than others. These outliers are scattered all along the chromosome and represent ‘hot spots’ of variability: genes whose fluctuation around the ‘ideal value’ (correspondent to their PC1 score) is very high. The peculiar relation of ‘hot-spots’ with their ‘cell-kind’ specific expression value is depicted in Fig.4.

Keeping in mind all the components have by construction zero mean and unit standard deviation on the whole set, it is evident that the same amount of total variability is constrained into 3 standard deviation interval for PC1 (see Fig.4), while fluctuation modes (PC2 and PC3) variance is driven by outliers (gene expression values outside the 3 sigma interval)); note that these outliers are leading contributors to average expression of PC2 and PC3 genes.. PC1 (cell-kind attractor) corresponds to the chromatin organization on the global scale (so giving rise to a continuous expression value distribution), fluctuation modes are driven by discrete local variations in flexibility correspondent to what we called ‘hot spots’. The results of both Fig.3 and Fig.4 are identical for HRG (data not shown). Table 2 reports the PCA solution for HRG condition;

**Table 1a.** Eigenvalues distribution (HRG)

| **Component** | **Eigenvalue** | **Explained Variance**<br>**(%)** | **Cumulative variance**<br>**(%)** |
| --- | --- | --- | --- |
| **PC1** | **17.63** | **97.95** | **97.95** |
| **PC2** | **0.129** | **0.72** | **98.67** |
| **PC3** | **0.065** | **0.36** | **99.04** |
| **PC4** | 0.040 | 0.22 | 99.26 |
| **PC5** | 0.031 | 0.17 | 99.43 |

**Table 1b.** Loading Pattern (HRG)

| **LABEL** | **TIME(min)** | **PC1** | **PC2** | **PC3** |
| --- | --- | --- | --- | --- |
| **T0** | 0 | 0,984 | 0,046 | -0,023 |
| **T10** | 10 | 0,983 | 0,150 | 0,034 |
| **T15** | 15 | 0,956 | 0,280 | 0,064 |
| **T20** | 20 | 0,992 | -0,004 | -0,089 |
| **T30** | 30 | 0,989 | 0,018 | -0,128 |
| **T45** | 45 | 0,993 | 0,000 | -0,076 |
| **T60** | 60 | 0,991 | 0,011 | -0,106 |
| **T90** | 90 | 0,994 | -0,025 | -0,040 |
| **T2H** | 120 | 0,995 | -0,020 | -0,006 |
| **T3H** | 180 | 0,994 | -0,022 | 0,017 |
| **T4H** | 240 | 0,992 | -0,071 | 0,050 |
| **T6H** | 360 | 0,995 | -0,046 | 0,038 |
| **T8H** | 480 | 0,994 | -0,055 | 0,053 |
| **T12H** | 720 | 0,994 | -0,042 | 0,040 |
| **T24H** | 1440 | 0,990 | -0,083 | 0,060 |
| **T36H** | 2160 | 0,994 | -0,044 | 0,034 |
| **T48H** | 2880 | 0,994 | -0,047 | 0,039 |
| **T72H** | 4320 | 0,990 | -0,035 | 0,040 |

Comparing Table 2 with Table 1, it is worth noting that, as expected, fluctuation components are more relevant in terms of explained variance with respect to EGF (PC2 goes from 0.41% to 0.72% of explained variance, PC3 goes from 0.26% to 0.36%). But, still more important, the third time point (15 min) is both the one less correlated with ideal profile (0.955) and where the most relevant fluctuation component (PC2) has the by far more relevant influence (loading = 0.28). This is consistent with the observation that time point 3 (15 min) shows a drastic decay in gene expression profiles correlation with respect to the initial time *t* = 0 condition (see Fig. 5, Tsuchiya, 2015) particularly evident in the super critical state.

**Figure 5:**
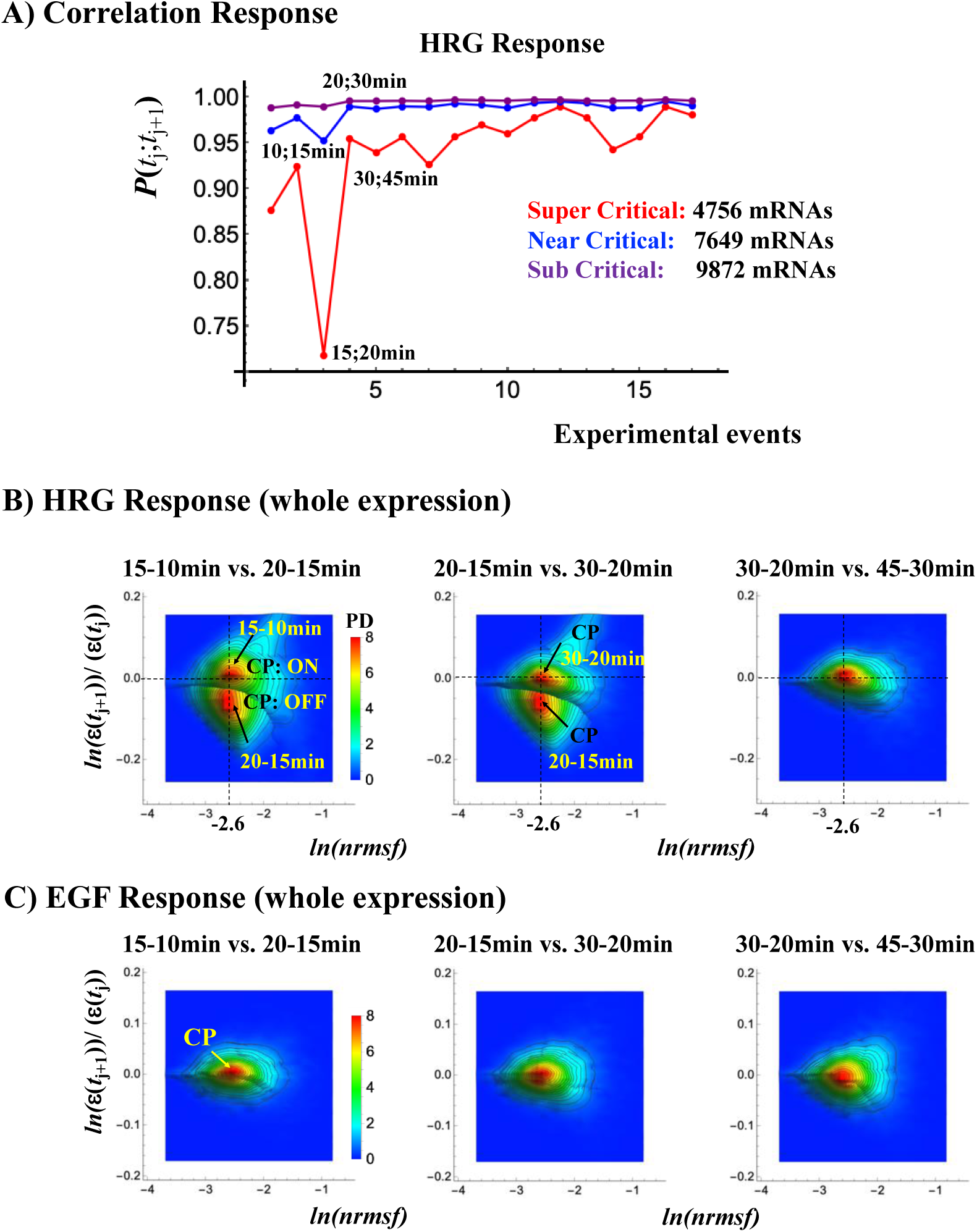
A singular response in critical states corresponds to the ON-OFF switch of critical point (CP). A) The dynamics of gene expression profile correlation between adjacent time points. At tipping point (15-20 min) there is a decay in correlation (maximal distance from stable profile) particularly evident in the super-critical state. **B)** Probability density functions (PDFs) of whole expression in HRG response is estimated in the space *nrmsf* (*x-*axis) vs. natural-log of fold change in expression (*y-*axis). The combined PDFs plot of adjacent time points shows the ON-OFF switch of the CP: ON at 10-15min and OFF at 15-20min, where the CP corresponds to the peak of probability density (red area). C) Same as B, for EGF response. The B) and C) reveal that, while in HRG response, a big avalanche in genome expression occurs, only local fluctuations are evident in EGF response (see main text and Supplementary file B).

The exceedingly high loading of the third time point (T3 = 15 minutes) on PC2 main oscillatory mode is consistent with the ON-OFF switch of the CP (see Figure5 and Supplementary File B) highlighted by previous SOC analysis (Tsuchiya, 2019).

The oscillation in time of the two main regulatory components is evident in Figure 6 reporting the loadings on PC2 and PC3 of the different time points. Loadings correspond to the relative importance of the two modes in determining the observed gene expression pattern at different times.

**Figure 6:**
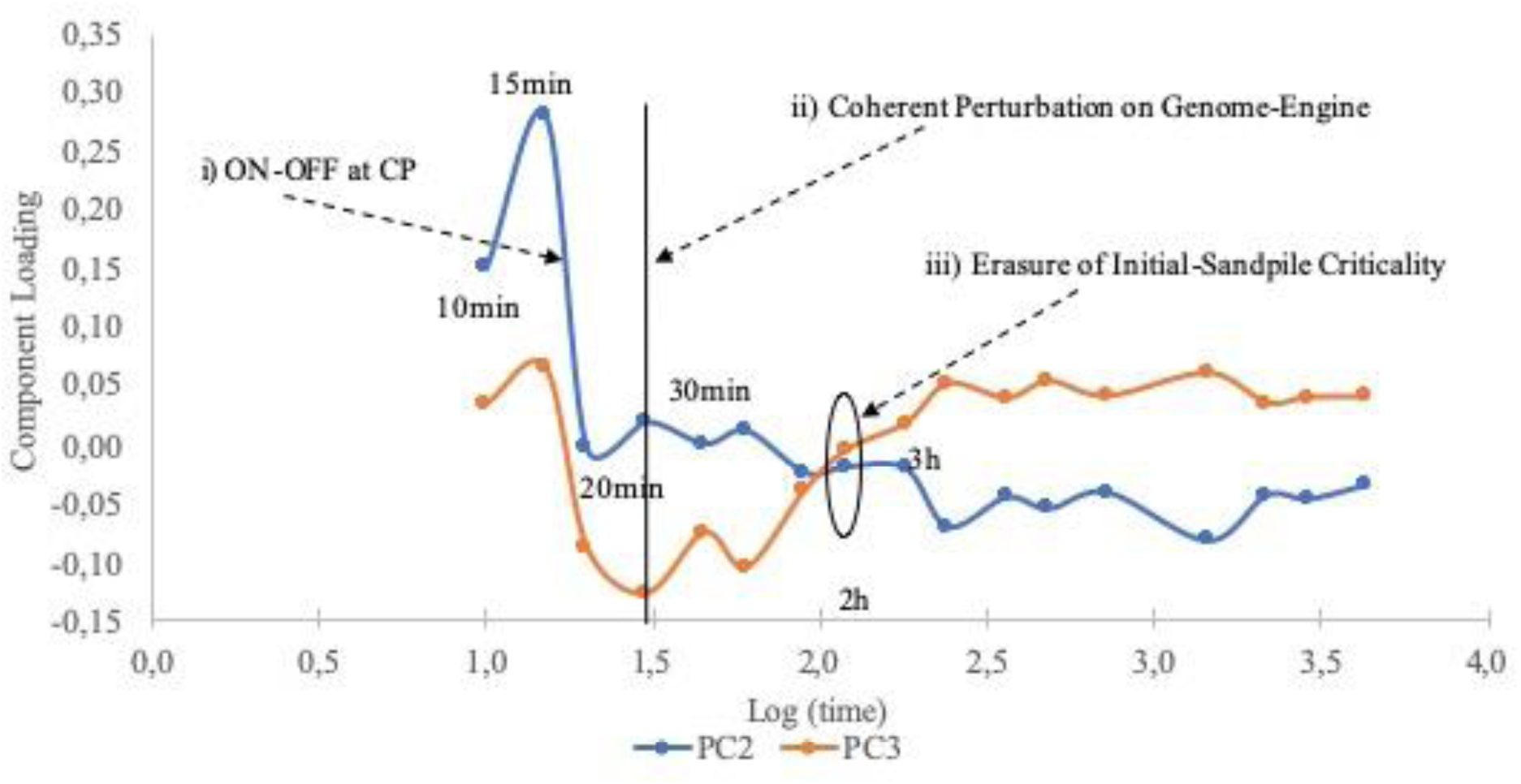
Temporal variation of regulatory modes (PC2, PC3) for HRG in relation to SOC events. PCA oscillation coincides with major SOC events: i) The ON(15min) - OFF(20min) at the CP (change in genome-attractor) induces a genome-wide avalanche, ii) after 30 min, the coherent perturbation on the genome-engine occurs for the preparation of the cell-fate change, and iii) initial sandpile CP disappears after 3h, suggesting the occurrence of the cell fate change. In general, the two modes behave as a sin/cosine pair (antiphase) in time so confirming their non-random character. This antiphase stems from the genome-engine, dominant expression flux between super (ensemble of high variance genes)- and sub (ensemble of low variance genes: majority from PC1 genes)-critical states

Going back to equation (3), we can easily appreciate how the oscillatory modes affect all the genes (with varying intensities), in fact these modes are not only responsible for cell fate transitions, but during ‘business as usual’ periods, they adapt the gene expression levels to the continuous microenvironment fluctuations. The ‘gene expression fluctuations’ in the normal condition are instrumental to keep largely invariant the cell-kind ideal profile. This is exactly the essence of Self Organized Complexity: the system is maintained in a constantly changing condition (correspondent to small avalanches in the sandpile model) around an ideal profile. Thus, PC2 and PC3 modes are the counterpart of the local regulation spread all along the genome we put in evidence by RQA in the previous chapter. This role is evident by the ability of PC1, PC2 and PC3 to predict the *nrmsf* of single genes (Table 3).

**Table 3a:** EGF condition.

| Component | *t-value* | **p** |
| --- | --- | --- |
| **PC1** | 45.74 | <0.0001 |
| **PC2** | -97.05 | <0.0001 |
| **PC3** | -21.31 | <0.0001 |

**Table 3b:** HRG condition.

| Component | *t-value* | **p** |
| --- | --- | --- |
| **PC1** | 58.24 | <0.0001 |
| **PC2** | -83.29 | <0.0001 |
| **PC3** | -61.29 | <0.0001 |

Table 3 gives a clear proof-of-concept of our model; global gene expression modes allow to reconstruct a single gene feature (its normalized mean square fluctuation: *nrmsf*) computed over the entire 18 time points by only three regressors (PC1, PC2, PC3). It is worth noting that minor components, explaining less than 1/1000 of PC1 variance, have an influence in the model (as expressed by *t*-values of the different regression coefficients) equal (or greater) than PC1. This is consistent with the fact that PC2 and PC3 correspond to relatively ‘free fluctuation’ of the system, while the motion along PC1 is much more constrained.

The physical counterpart of this statistical evidence is that variation along PC1 corresponds to shift the ‘center of mass’ (CM) of the system that can lead jumping off the attractor (transition). If this is the case, we expect that the genes with near zero values of PC2 and PC3 are exactly the ‘genome vehicle’ (Tsuchiya, 2010) those genes that cause the critical behavior of the SOC regulated system.

This genome vehicle can be seen from our current SOC result (Tsuchiya, 2010) in that the CP (critical point) acts as genome-attractor, which represents a specific set of critical genes with an activated (ON) or deactivated (OFF) state (see SOC summary in Introduction). In HRG, the CP is ON at 15 min and then OFF at 20 min through the coil-globule transition at the CP (Figure S2 in Supplementary file B). This ON-OFF switch of the CP state at 15-20min, i.e., change in the genome-attractor induces a big avalanche effect on genome expression. This is revealed (Fig.6) by the decay of PC2 and PC3 response at 15-20min.

It is important to compare EGF and HRG conditions, from here onward PC1, PC2 and PC3 of the EGF and HRG will be indicated as EGF1, EGF2, EGF3 and HRG1, HRG2, HRG3 respectively.

Both EGF and HRG mediated perturbations operate on the same cell-kind (MCF7 cells), if our interpretation of the meaning of different components is correct, we should find a complete superposition between the first components of EGF and HRG (EGF1 and HRG1 respectively) given they refer to the same cell-kind ideal profile. That is exactly what we find with a near to unity correlation between EGF1 and HRG1 (left top panel of Figure 7).

**Figure 7:**
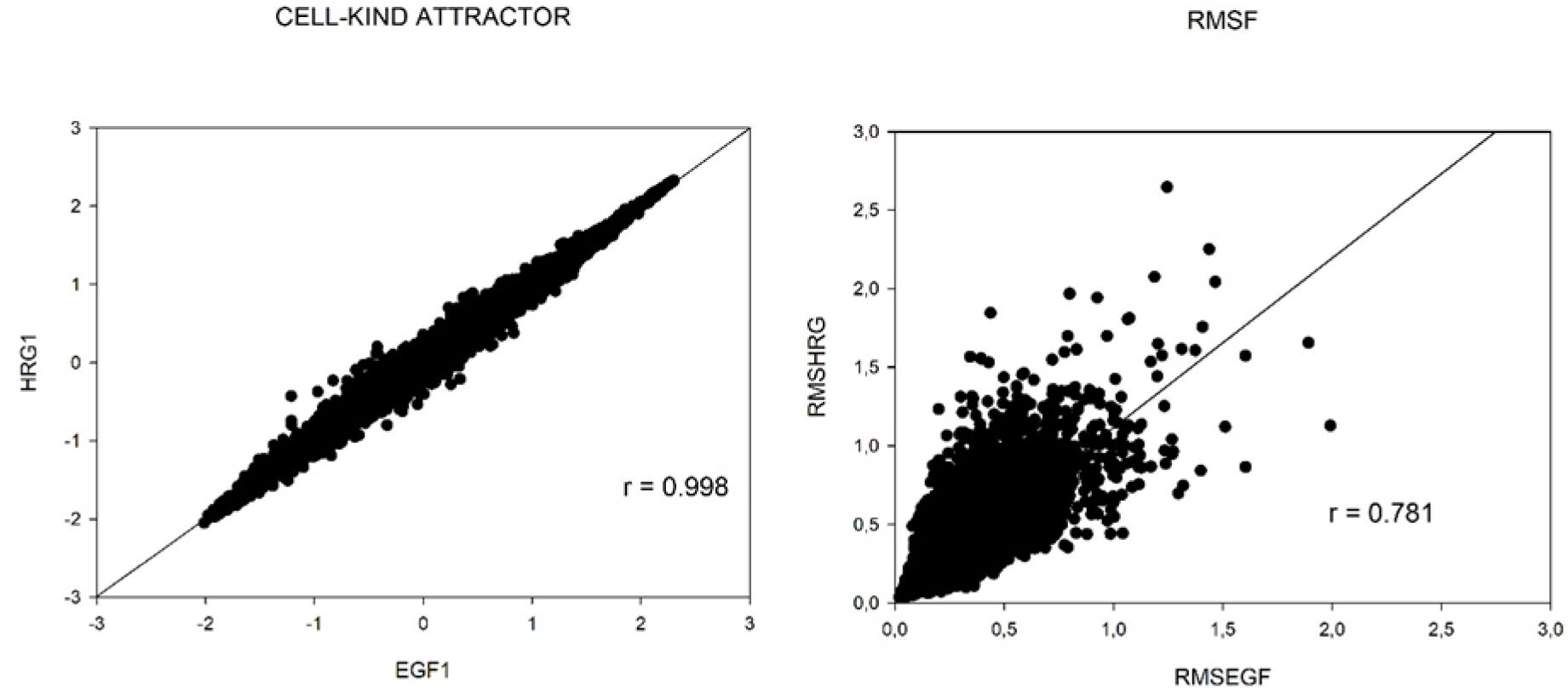
EGF and HRG comparison. The figure reports two different comparisons between EGF and HRG cases: A) Left panel has as axis the first component of the two conditions (HRG1 and EGF1 respectively). The near to unity correlation of HRG1 and EGF1 scores demonstrates that the first component corresponds to the cell-kind (shared by the two conditions). B) Right panel shows the single genes fluctuation in time (rmsf) correlates between the two EGF and HRG cases, pointing to the fact that ‘fluctuation entity’ is an intrinsic property of each gene largely independent of the nature of the stimulus.

Shifting to *nrmsf* (entity of fluctuation in time of genes), the correlation between EGF (*x* axis) and HRG (*y* axis) scores a correlation coefficient *r* = 0.78, (top right panel of Figure 7). This relation is statistically very significant and implies that the two EGF and HRG stimuli impinge over a highly superimposable set of genes. This favors the mainly positional character of gene expression regulation (see paragraph 3.3).

The HRG and EGF comparison encompasses some useful tracks to understand the difference between a stimulus that is able to induce a transition (HRG) and one that is not (EGF).

The first ‘track’ is the observation that HRG *nrmsf* is (on average) greater than EGF one (0.28 vs. 0.22) points to a greater applied energy in the transition case. Indeed, any detachment from ideal expression value can be considered as provoked by the need to dissipate an excess of energy, being the cell-kind attractor located in an energy minimum. The second track was suggested by the entrainment into a resonant peak of HRG2 and HRG3 depicted in Fig.5 and Fig.6, this resonant peak is absent in EGF case (see Fig.5 Panel C), this resonance drastically increases the applied energy on the genome.

These results are consistent with SOC analysis in that ON-OFF switch of the CP (i.e., provoking genome avalanche through change in the genome-attractor) occurs at 15-20min through transitional change in singular behaviors at the CP for HRG cell response, whereas the CP is at OFF state in EGF cell response (i.e., no genome avalanche).

The analysis of log-normalized *rmsf*:

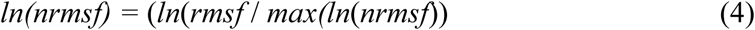

allows to appreciate the convergence of the SOC (more theoretical) and PCA (more empirical) approaches, this parameter is the order parameter of self-organization (Tsuchiya, 2015-2106, see Supplementary file B).

According to SOC, we do expect a value within −2.5∼-2.6: singular behaviors of the CP (see Figure S2.2-3 in Supplementary file B): it occurs consistently around −2.58 for both HRG and EGF.

Figure 8 reports the space spanned by EGF2 and log-normalized *rmsf* on the top and the 3D space with added EGF3 (bottom panel).

**Figure 8:**
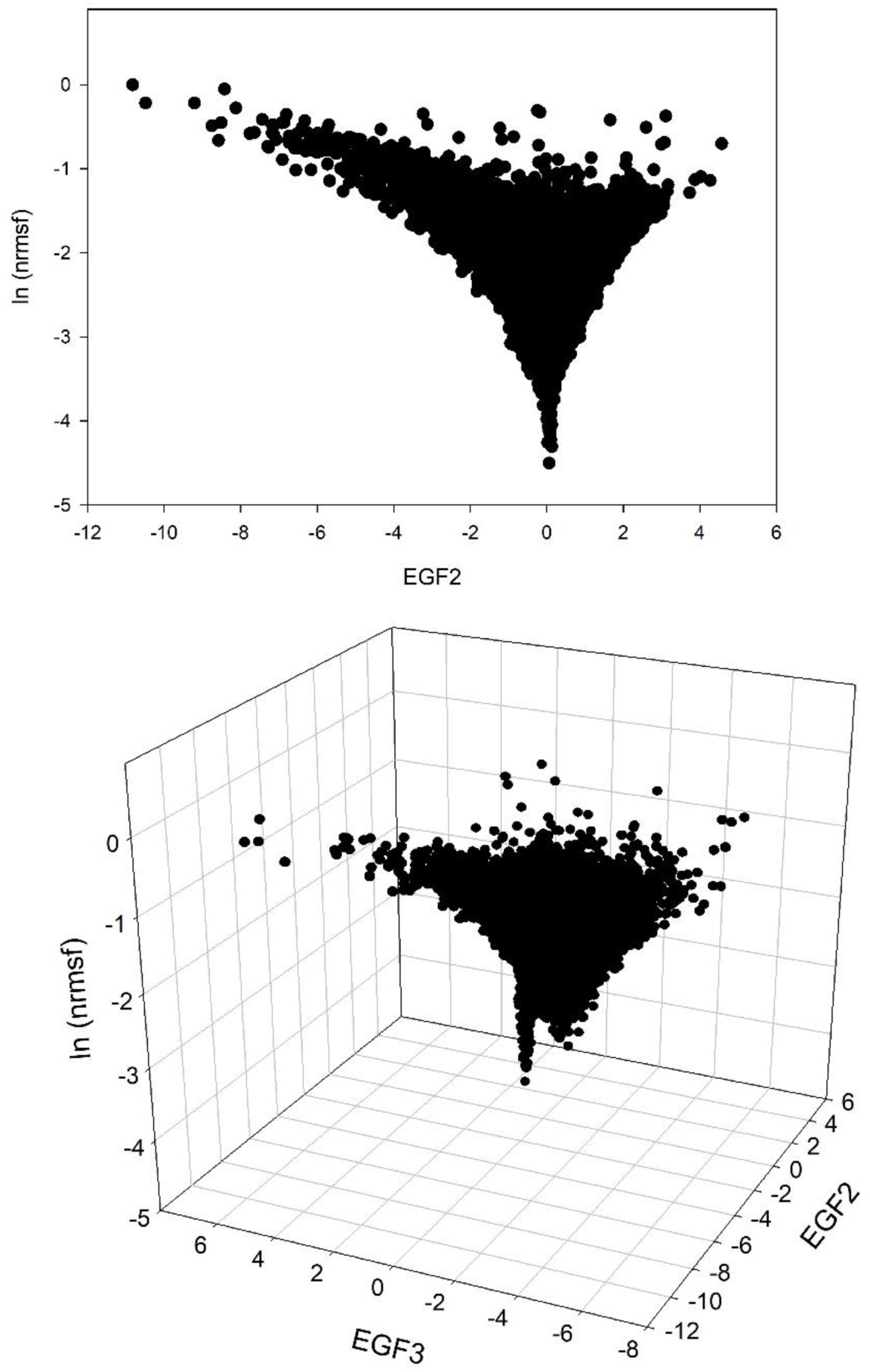
SOC critical parameter in terms of oscillating expression modes (EGF, no transition case). The bottom of the funnel corresponds to the ‘core’ of the attractor (CP, genes with very low fluctuations that consequently have near zero contribution of fluctuations modes (EGF2, EGF3).

Figure 9, which is relative to the HRG transition case, it is very similar to the EGF case but with some subtle differences. The presence of a clear funnel landscape with a minimum of fluctuation (near zero PC2) mode at very low *ln*(*nrmsf*) is evident in both EGF and HRG cases. This is consistent with the fact that zero values of oscillating modes correspond to the ‘invariant core’ (genome-attractor) of cell-kind profile and suggests that CP genes (low values of *ln*(*nrmsf*)) correspond to those invariant (and thus zero valued as for oscillating modes). In EGF case (Figure 8) there is a unimodal minimum exactly located at *t*= 0 (minimum influence of PC2). On the contrary, in the case of HRG (top panel of Fig.9) the minimum of the main oscillating mode (HRG2) is splitted in a bimodal way around zero. In figure 5B, the second peak in the HRG response has to do with the signature of ON-OFF transition of the CP.

**Figure 9:**
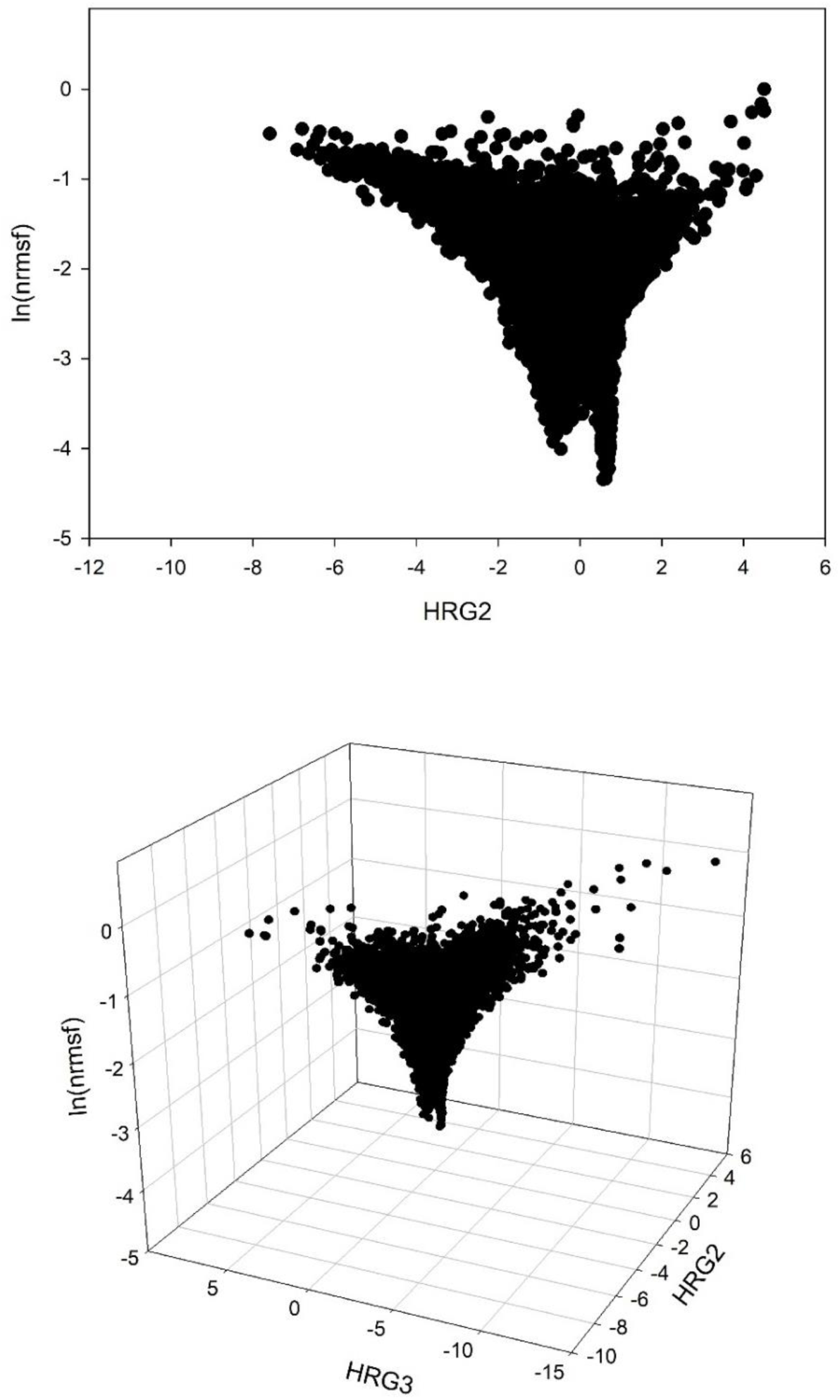
SOC critical parameter in terms of oscillating expression modes (HRG, transition case). The bottom of the funnel corresponds to the ‘core’ of the attractor (CP, genes with very low fluctuations that consequently have near zero contribution of fluctuations modes (HRG2, HRG3).

The main proof of the convergence between the theoretical (SOC) and bottom-up (PCA) approach is given by the possibility to estimate the SOC-theoretically predicted values of *ln*(*nrmsf*) by means of principal component scores (Figure 10).

**Figure 10:**
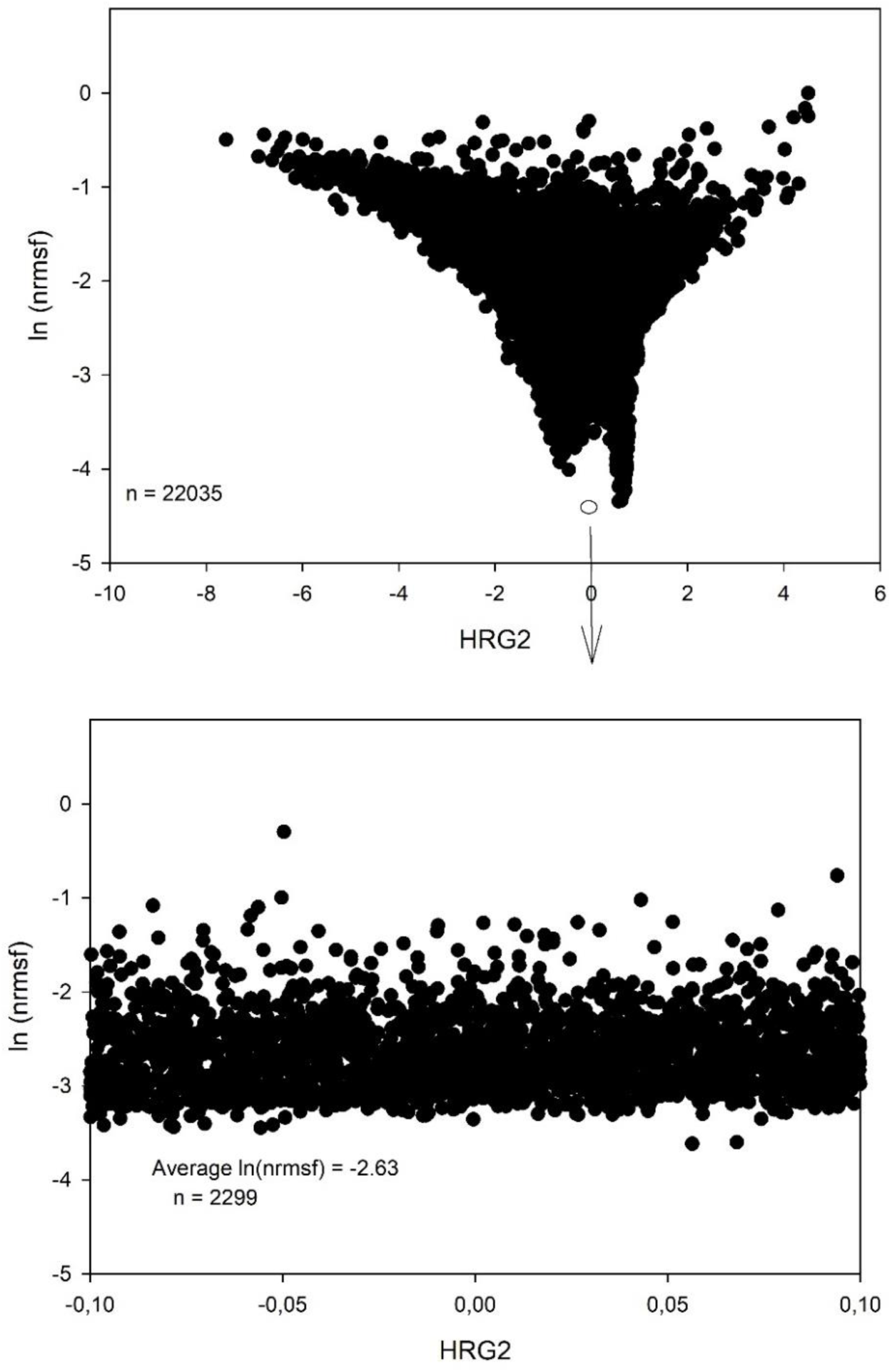
The core of cell-kind attractor corresponds to CP. The figure depicts, in the HRG transition case, the ability of second component (main oscillating mode) to exactly predict the theoretical value of Critical Point (CP) in ln(*nrmsf*) terms. The top panel reports the bi-dimensional space spanned by HRG2 and ln(*nrmsf*) with the evidence of the bimodal negative peak correspondent to near zero values of HRG2. Bottom panel reports the estimation of ln(*nrmsf*) in terms of average over 2299 genes in the vicinity of zero HRG2 score (from −0.1 to 0.1 standard deviation units).

Top panel of Figure 10 reports the bimodal minimum of *ln*(*nrmsf*) for near 0 values of HRG2. Bottom panel is an inset focusing on HRG2 scores between −0.10 and +0.10. This range corresponds approximately to 1/10 of total number of genes, the average *ln*(*nrmsf*) of these genes is −2.63, which is similar to the estimation from SOC, *ln*(*nrmsf*) = −2.58 relative to the CP obtained by a completely different paradigm.

### 3.3 Functional and positional regulation

A basic tenet of functional genomics is that the genes most affected by an external perturbation (being it a disease, a drug, a toxicant, etc.) constitute a functional signature of the perturbation or, in other words, the ‘molecular mechanism’ allowing the perturbation to exert its effect. Crow et al. put the above ‘dogma’ in crisis in a recent work (Crow, 2019).

The authors performed an accurate meta-analysis involving over 600 differential gene expression studies from the Gemma database (https://gemma.msl.ubc.ca) involving human cell lines from different tissues and conditions spanning from the effect of mutations to drug and disease response. They put together the differentially expressed (DE) sets of genes from the 600 studies in order to generate a predictor based on the 200 genes most frequently observed in the different DE sets. This empirical DE set can be considered (due to the heterogeneity of conditions from which it derives) as a signature largely independent from specific functional features (biological mechanisms). This empirical ‘prior probability’ based signature performs very well in discriminating different experimental conditions (mean area under the Receiver Operating Characteristic curve, ∼0.8 (ROC)), indicating that a large fraction of DE hit lists is nonspecific. In contrast, predictors based on attributes such as gene function, mutation rates, or network features perform poorly (Crow, 2019). Genes associated with sex, the extracellular matrix, the immune system, and stress responses are prominent within the “DE prior”. The authors demonstrated that increasing the ‘specificity’ of the DE does not increase the discrimination power. The authors correctly explain their revolutionary results in terms of the redundancy of biological regulation taking place downward the DNA transcription level.

In summary they state that, given the biological systems have fully connected regulation networks at different hierarchical levels from DNA transcription to metabolism, an initially largely functionally not-specific stimulus can give rise to a specific response in the subsequent cascade of ‘processing’ of the initial information by biological system. This allows very subtle initial differences between DE gene sets to provoke differential (function specific) biological consequences.

The Crow et al. results are in line with our results:

1. RQA tells us that gene co-regulation happens on a purely positional (not functional) basis.
2. PCA allowed very similar regulation components to emerge from stimuli having different biological consequences on the system at hand.
3. SOC is a statistical mechanics phenomenon blind to gene specific function. In order to empirically connect our study to Crow et al. ‘DE prior’ (generic signature) we computed the correlation between the relevant genes selected from our study (‘hot spots’ (genes more affected by regulatory oscillation modes) and CP (critical point genes responsible for global genomic level avalanche)) with the functional spectrum of ‘DE prior’ (generic signature).

In order to perform this task, we generated by means of PANTHER Gene Ontology software (http://pantherdb.org/) the ‘molecular function’ and ‘biological process’ profiles of ‘hot-spots’ and the CP lists for both EGF and HRG conditions and compared these profiles with the DE prior (Generic) gene list (Crow, 2019).

We generated lists of 200 genes by three selection criteria: a) Maximal PC2 scores, b) minimal PC2 scores (both (a) and (b) selection criteria point to hot spots) and c) Critical Point (CP) genes (near zero values of PC2). These criteria were applied to both EGF and HRG conditions so generating six different lists of genes. The ‘molecular function’ and ‘biological process’ profiles relative to these six lists were correlated with the corresponding profile of 200 genes list of the Crow et al. ‘DE prior’ (Generic). Figure 11 below reports the striking correlation between our lists and the ‘DE prior’ list:

**Figure 11:**
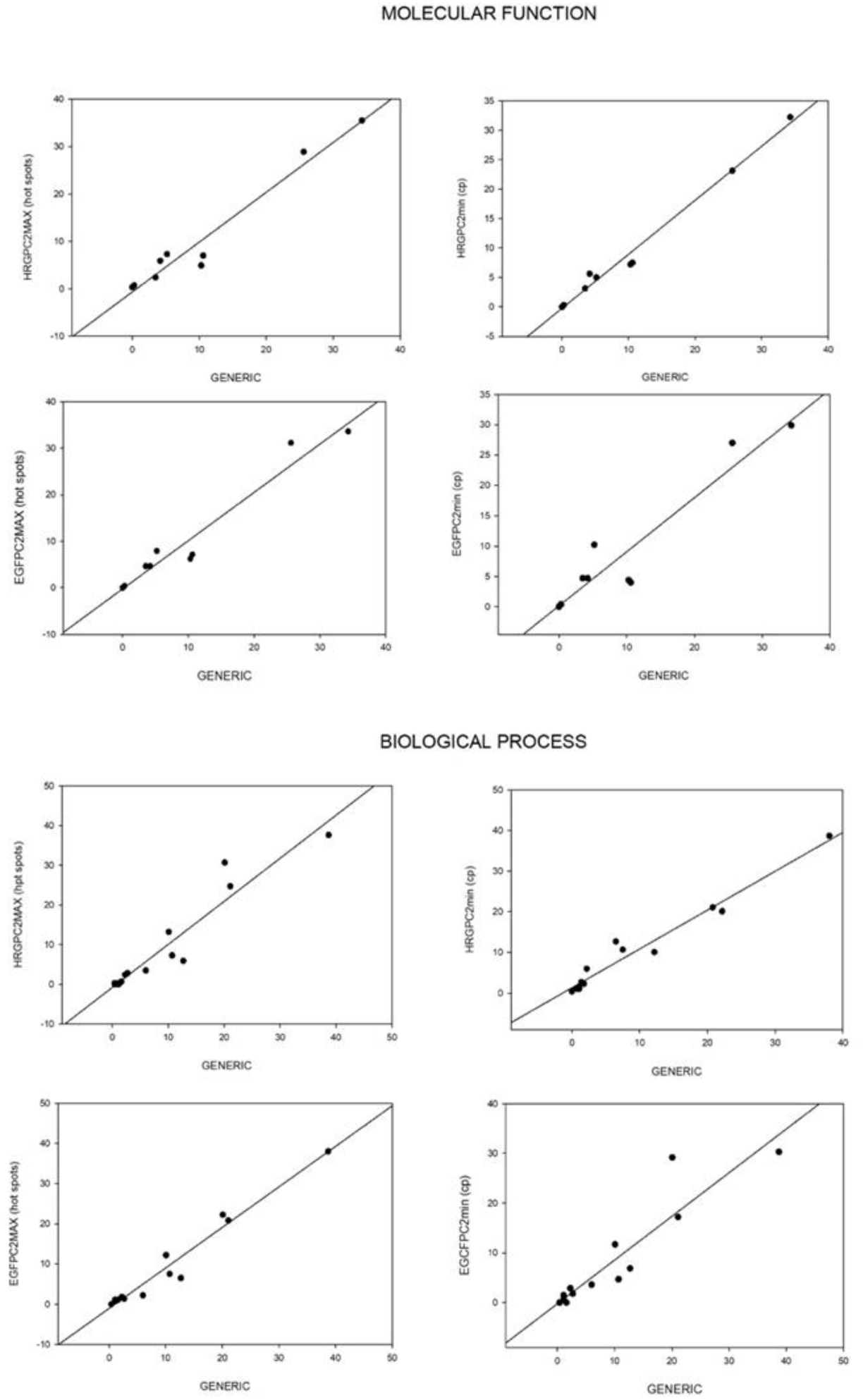
Correlation between ‘hot-spots’ and CP spectra with Generic signature (DE prior). Vector points correspond to molecular function (9 values) and biological process (14 values) categories, all the correlation (Pearson *r*) were statistically significant and with *r*> 0.90 pointing to a striking superposition between meta-analysis derived generic signature and genes relevant for SOC regulation.

The observed superposition between our and Crow et al. results gives a physical plausibility to the observed specificity of biological regulation without relying on improbable ‘Maxwell’s demons’ able to select the ‘right genes’ interspersed in a 2 meters long molecule constrained in the few microns space of cell nucleus by a drastic folding compression.

## Discussion and Conclusion

In this work, by means of data-driven techniques (PCA and RQA), we validated the role of chromatin remodeling as the material basis of the Self-Organized Critical Control (SOC) of genome expression. The critical point (CP) acts as the genome-attractor, which dynamically self-organizes whole expression consequently, drastic changes in the CP induce genome-wide expression avalanche. RQA and PCA demonstrated this avalanche stems from coordinated activity of positional-based (not biological functional) local chromatin interaction. The hot spots are the generators of genome-engine revealed by the SOC model. Furthermore, in HRG response, the activation of the CP entrains hot spots (genes) in a resonance mode (critical transition), but not in EGF response. Expression of the CP changes in time for both HRG and EGF responses albeit invariant temporal expression variance (*nrmsf*) of the CP, suggesting that there exists threshold for its ON-OFF state, i.e., the activation mechanism of the CP. This elucidation is expected to provide a novel cell-fate control mechanism.

In chromatin remodeling perspective, we summarize a cell-fate transition by this s cell response:

1. Establishing a new ‘preliminary’ cell-fate: HRG activates the ErbB receptor with sustained extracellular signal regulated kinase (ERK) activity (Nagashima, 2007). This induces a sufficiently strong perturbation (mirrored by a peak of PC2 and PC3 explained variance) on the MCF cell genome, i.e., setting the CP to ON by a consequent shift to swollen coil state before 15min. The activation of the CP genes occurs through a purely positional (not functional) basis of chromatin. This ON state of the CP leads to a new ‘preliminary’ cell-fate accompanying with the bifurcation of the super-critical attractor at 15min (Figure 5).
2. Setting the CP to OFF shifting to compact globule state: A critical transition occurs at the CP from the swollen-coiled state to compact-globule state, which sets the CP from ON to the OFF state during 15-20min. This exactly concedes with the timing of the resonant mode of PC2 and PC3 (Figure 5,6) at 10-20min. This released energy ignites the local and positional chromatin to coherent first order transition on chromatin (genome-wide expression avalanche).
3. Switch of coherent perturbation on the genome-engine: The perturbation on the CP (the genome-attractor) induces the switch of coherent perturbation on the genome-engine from suppression to enhancement on the dominant cyclic expression flux between super- and sub-critical state attractors (Supplementary file B). This coherent perturbation drives the cell-fate decision (Tsuchiya, 2019)
4. Stabilization of a new cell-fate: After 20min, gradual stabilization (Figure 5,6) of a new cell-fate occurs by positional chromatin remodeling, which is consistent with the SOC event that implies the apparent conundrum (see kinetic energy flux in Supplementary file B) of a dynamically stable critical state.
5. Attainment of a new cell-fate attractor: After 3h, the initial-state criticality is erased (i.e., erasure of the memory of the initial-state CP, Tsuchiya, 2016) which indicates that the genome system attains a new cell-fate attractor (Figure 5,6).
6. Sub-critical vs. super-critical in the genomic state: In EGF-stimulated response, EGF-stimuli do not elicit the CP ON state and the perturbation is locally confined. Thus, the genome state (at cell-population level) of EGF-stimulated system, at odds with HRG condition, remains ‘sub-critical’ (no cell-fate change).

It is worth noting the universality of SOC mechanism in biology: the motion along PC2 and PC3 are orthogonal with respect to PC1 thus do not affect the ‘center of mass’ (CM) of the system, on the contrary their oscillating behavior in time, allows for maintaining the system on its ‘attractor’ profile dissipating external perturbations. This is exactly what happens in molecular dynamics with the fluctuation of amino-acid residues keeping largely invariant the 3D structure of the system, it is not a case that proteins are SOC governed systems too (Phillips, 2009). The unique property of SOC of being a ‘critical attractor state’ allows proteins to manage with thermal motion without losing their global 3D configuration while in the same time (thanks to CP-like amino-acid residues) responding with fast transition to allosteric stimuli.

The time-dependent behavior of the genomic DNA transition follows a kinetic equation that shows cubic nonlinearity (Takagi, 1999). Cubic-type nonlinearity plays an essential role in nervous system excitability (see, for example, the FitzHugh–Nagumo-type equation to show cubic nonlinearity for the fundamental characteristics nerve firing, FitzHugh, 1995; Izhikevich, 2006). Furthermore, SOC is at play in neuronal networks (Plenz, 2014; Shew, 2009), which indicates a non-trivial similarity between the coherent network of genomic DNA transitions and neural networks.

Overall, we demonstrate the feasibility of a ‘biological statistical mechanics’ spanning different organization layers of biological material that can push the research toward a more integrative and physically motivated view of biology.

## Supporting information

Essential summary of SOC results and RQA method

## Acknowledgments

We thank Prof. Kenichi Yoshikawa for his continuous interest and inspiration.

## References

Bohn, M., Diesinger, P., Kaufmann, R., Weiland, Y., Müller, P., Gunkel, M., Cremer, C. (2010). Localization microscopy reveals expression-dependent parameters of chromatin nanostructure. Biophysical journal, 99(5), 1358–1367. doi:10.1016/j.bpj.2010.05.043

Cattell, R. B. (1966). The scree test for the number of factors. Multivariate Behavioral Research, 1(2), 245-276

Chang, H. H., Oh, P. Y., Ingber, D. E., & Huang, S. (2006). Multistable and multistep dynamics in neutrophil differentiation. BMC cell biology, 7(1), 11

Cremer, T., Küpper, K., Dietzel, S., & Fakan, S. (2004). Higher order chromatin architecture in the cell nucleus: on the way from structure to function. Biology of the Cell, 96(8), 555–567

Crow, M., Lim, N., Ballouz, S., Pavlidis, P., & Gillis, J. (2019). Predictability of human differential gene expression. Proceedings of the National Academy of Sciences, 116(13), 6491-6500

FitzHugh R (1955) Mathematical models of threshold phenomena in the nerve membrane. Bull Math Biophys 17: 257–278

Giuliani, A., Piccirillo, G., Marigliano, V., & Colosimo, A. (1998). A nonlinear explanation of aging-induced changes in heartbeat dynamics. American Journal of Physiology-Heart and Circulatory Physiology, 275(4), H1455-H1461

Giuliani, A., Tsuchiya, M., Yoshikawa, K. (2018). Self-Organization of Genome Expression from Embryo to Terminal Cell Fate: Single-Cell Statistical Mechanics of Biological Regulation. Entropy 20, 13

Izhikevich, E. M., and R. FitzHugh. 2006. FitzHugh-Nagumo model. Scholarpedia 1:1349

Jolicoeur, P., & Mosimann, J. E. (1960). Size and shape variation in the painted turtle. A principal component analysis. Growth, 24(4), 339-354.

MacArthur, B. D., Ma’ayan A, Lemischka IR. (2009) Toward stem cell systems biology: from molecules to networks and landscapes. Cold Spring Harb Symp Quant Biol.73:211–5. doi: 10.1101/sqb.2008.73.061

Mayer, R., Brero, A., Von Hase, J., Schroeder, T., Cremer, T., & Dietzel, S. (2005). Common themes and cell type specific variations of higher order chromatin arrangements in the mouse. BMC cell biology, 6(1), 44

Marwan, N., Romano, M. C., Thiel, M., & Kurths, J. (2007). Recurrence plots for the analysis of complex systems. Physics reports, 438(5-6), 237-329

Nagashima, T., Shimodaira, H., Ide, K., Nakakuki, T., Tani, Y., Takahashi, K., Yumoto, N., Hatakeyama, M. (2007) Quantitative transcriptional control of ErbB receptor signaling undergoes graded to biphasic response for cell differentiation. J Biol Chem 282: 4045–4056

Orlando, G., Zimatore, G., (2017). RQA correlations on real business cycles time series Pramana - Journal of Physics - Indian Academy of Sciences Conference Series 1(1), DOI:10.29195/iascs.01.01.0009

Phillips, J. C. (2009) Scaling and self-organized criticality in proteins: Lysozyme c. Physical review E 80.5: 051916

Plenz D, Niebur E (2014) Criticality in neural systems. Wiley-VCH, Germany

Shew WL, Yang H, Petermann T, Roy R, Plenz D (2009) Neuronal avalanches imply maximum dynamic range in cortical networks at criticality. J Neurosci 29(49): 15595–15600

Raser, J. M., O’Shea, E. K., (2005). Noise in gene expression: origins, consequences, and control. Science, 309(5743):2010–3

Saeki, Y., Endo, T., Ide, K., Nagashima, T, Yumoto, N., Toyoda, T., Suzuki, H., Hayashizaki, Y., Sakaki, Y., Okada-Hatakeyama M. (2009). Ligand-specific sequential regulation of transcription factors for differentiation of MCF-7 cells. BMC Genomics 10: 545

Takagi S, Yoshikawa K (1999) Stepwise collapse of polyelectrolyte chains entrapped in a finite space as predicted by theoretical considerations. Langmuir 15: 4143–4146

Tsuchiya, M., Vincent, P., Giuliani Tomita, M., Kumar Selvarajoo, K. (2010). Collective Dynamics of Specific Gene Ensembles Crucial for Neutrophil Differentiation: The Existence of Genome Vehicles Revealed, PLoS One 5, e12116

Tsuchiya, M., Hashimoto, M., Takenaka, Y., Motoike, I. N., Yoshikawa, K. (2014). Global genetic response in a cancer cell: Self-organized coherent expression dynamics. PLOS One 9: e97411

Tsuchiya, M., Giuliani, A., Hashimoto, M., Erenpreisa, J., Yoshikawa, K. (2015). Emergent Self-Organized Criticality in gene expression dynamics: Temporal development of global phase transition revealed in a cancer cell line. PLoS One 11, e0128565

Tsuchiya, M., Giuliani, A., Hashimoto, M., Erenpreisa, J., Yoshikawa, K. (2016). Self-organizing global gene expression regulated through criticality: Mechanism of the cell-fate change. PLoS ONE 11: e0167912

Tsuchiya, M., Giuliani, A., Yoshikawa, K. (2019). Underlying Genomic Mechanism for Cell-Fate Change from Embryo to Cancer Development. Preprint, bioRxiv: doi: https://doi.org/10.1101/637033

van der Knaap J.A. and Verrijzer C.P. (2016) Undercover: gene control by metabolites and metabolic enzymes, Genes Dev. 30:2345–2369

Vickaryous, M. K., & Hall, B. K. (2006). Human cell type diversity, evolution, development, and classification with special reference to cells derived from the neural crest. Biological reviews, 81(3), 425–455

Webber Jr, C. L., & Zbilut, J. P. (1994). Dynamical assessment of physiological systems and states using recurrence plot strategies. Journal of Applied Physiology, 76(2), 965-973

Wolynes, P. G. (2010). Gene regulation: Single-molecule chemical physics in a natural context. In Single Molecule Spectroscopy in Chemistry, Physics and Biology (pp. 553–560). Springer, Berlin, Heidelberg

Yoshikawa, K., Takahashi, M., Vasilevskaya, V.V., Khokhlov, A.R. (1996). Large Discrete Transition in a Single DNA Molecule Appears Continuous in the Ensemble. Phys. Rev. Lett., 76, 3029–3031; 10.1103/PhysRevLett.76.3029

Yoshikawa K, Noguchi H (1999) A working hypothesis on the mechanism of molecular machinery, Chem Phys Lett 303: 10–14

Yoshikawa, K., Yoshikawa, Y. (2002). Compaction and Condensation of DNA, in Pharmaceutical Perspective of Nucleic Acid-Base Therapy. Tayler & Francis, Abingdon, UK, 137–163; 40

Zentgraf, H., & Franke, W. W. (1984). Differences of supranucleosomal organization in different kinds of chromatin: cell type-specific globular subunits containing different numbers of nucleosomes. The Journal of cell biology, 99(1), 272–286

Zinchenko, A., Pyshkina, O., Lezov, A., Sergeyev, V., Yoshikawa, K. (2008). Single DNA molecules: compaction and decompaction (chapter 3), in DNA interactions with polymers and surfactants. Wiley-Blackwell

Zimatore, G., Fetoni, A., Paludetti, G., Cavagnaro, M., Podda, M.V., Troiani, D., (2011) “Post-processing analysis of transient-evoked otoacoustic emissions to detect 4 kHz-notch hearing impairment – a pilot study,” Med Sci Monit 17(6): MT41–49.

Zimatore, G., Cavagnaro, M., (2015). “Recurrence Analysis of Otoacoustic Emissions,”. In Recurrence Quantification Analysis: Theory and Best Practices. 253–278. Springer, Cham.

Zimatore, G., Garilli, G., Poscolieri, M., Rafanelli, C., Gizzi, F. T., Lazzari, M., (2017) “The remarkable coherence between two Italian far away recording stations points to a role of acoustic emissions from crustal rocks for earthquake analysis,” Chaos: An Interdisciplinary Journal of Nonlinear Science, 27(4), 043101.

Zimatore, G, Orlando, G., (2018). “Recurrence quantification analysis of business cycles,” Chaos, Solitons and Fractals 110, 82–94 doi: 10.1016/j.chaos.2018.02.032

