## Supplementary material for "Self-Organization of Whole Gene Expression through Coordinated Chromatin Structural Transition: Validation of Self-Organized Critical Control of Genome Expression": Essential summary of SOC results and RQA method

### *Appendix*

#### 1 Appendix A: The self-organized criticality (SOC)

In physics, the self-organized criticality (SOC) is a property of dynamical systems that have a critical point as an attractor. SOC governed systems display the spatial and/or temporal scale-invariance characteristic of the critical point of a phase transition, but without the need to tune control parameters to a precise value, because the system spontaneously tunes to it as it evolves towards a critical state. The concept was put forward by Bak, Tang and Wiesenfeld (Bak, 1987,1988), and is considered one of the mechanisms by which complexity arises in nature. SOC arises in slowly driven non-equilibrium systems with a large number of degrees of freedom and strongly nonlinear dynamics. We define 'slowly-driven', those system that are not drastically perturbed by an external force but experience very mild but continuous external stimuli (this will be clear in the following when we will describe the so-called sand-pile model). This condition can be equated to the 'normal existence' of biological systems like tissues or cells in culture exposed to a continuously varying (but dynamically stable) microenvironment.

In the general definition of SOC we can recognise an (apparent) conundrum: how a critical (i.e. unstable) state can be at the same time an attractor (i.e. stable state)? To solve this conundrum, we need to introduce the notion of 1/f (or power spectrum) scaling.

##### 1.1 Supplementary Figure Appendix A

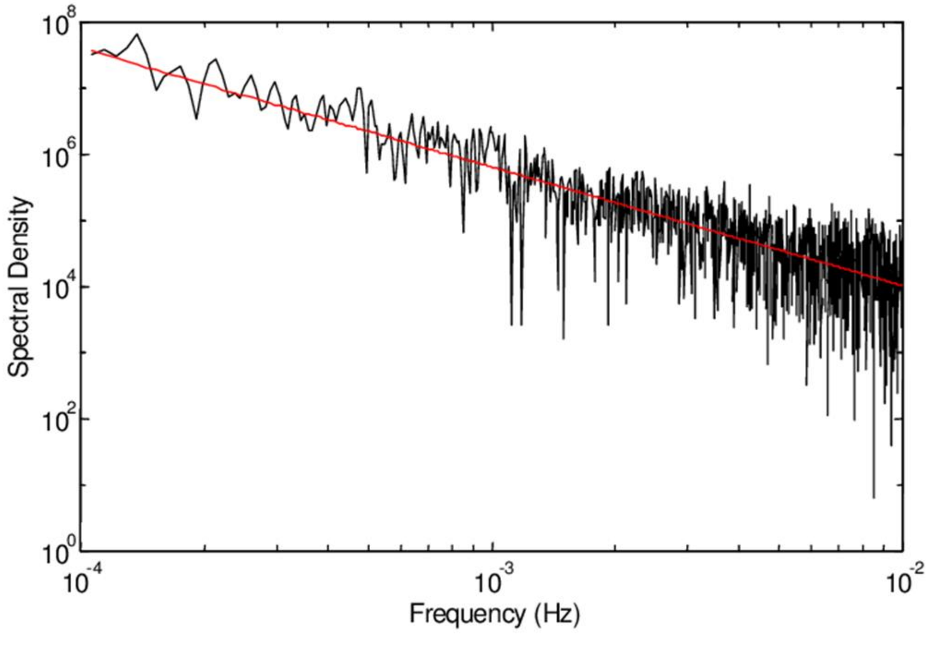


**Figure S1.1.** Pink noise (or 1⁄f noise) is a signal or process with a frequency spectrum such that the power spectral density (energy or power per frequency interval) is inversely proportional to the frequency of the signal. Pink noise is the most common kind of signal in biology and, in general, in all those phenomena keeping together processes and structures with a large range of spatial and time scales.

A classical paradigm of this organisation is the airports' network, with very few international hubs hosting a great number of flights like Frankfurt or New York and many small airports with a limited number of flights. This kind of distribution provokes a very peculiar statistical consequence. The asymmetric distribution of 1/f scaling does not allow the convergence to a "characteristic scale" of the system (scale free behaviour). In other words, if we get finite samples (any empirical sample is by definition finite even if it has a very high number N of elements) we will insert into the statistics the 'exceptionally big events' only at very high N. This happens for the simple reason these exceptionally big events are extremely rare: in the case of Gaussian distribution, the scarcity of 'exceptionally big events' is exactly balanced by the scarcity of 'exceptionally small events'. Thus, in the case of a feature with a normal distribution like the height of human beings (centred around 170-175 cm), if we increase the sampling, the uncertainty of the exact value of the population mean decreases. The exceedingly short and exceedingly tall persons balance their relative effects (law of large numbers) and the sample mean converges toward the population mean (characteristic scale) at increasing sample size. On the contrary, when this 'balance of extremes' does not happen (like in the 1/f scaling), at increasing sample size, the computed average is continuously shifted on the right and does not converge to a characteristic scale.

This is much more than a statistical curiosity if we keep in mind that the 1/f distribution refers to real biological systems and has a crucial role in SOC.

The paradigmatic SOC-based system is the sandpile: think of pouring sand very slowly (ideally one grain at a time), onto a flat, circular surface (this is exactly what we intend for 'slowly-driven').

At first, the grains stay close to the position where they land, very soon they start to accumulate, creating a pile that has a gentle slope. Going on with the experiment, when the slope becomes too steep, somewhere on the pile, the grains slide down, causing a small avalanche. As we add more sand, the slope of the pile further steepens, and the average size of avalanches increases (the size of avalanches follows a 1/f scaling: many small avalanches, very few huge avalanches).

The pile stops growing when the amount of sand added balances the amount of sand falling off the edge of the circular surface. At that point, the system reaches the critical state. This state is (dynamically) stable like any proper attractor: the continuous avalanches are counter-balanced by the added sand and the height and shape of the pile remains the same. Nevertheless, occasionally (right part of the 1/f spectrum), an added grain can cause a large catastrophic avalanche by a sort of chain reaction involving progressive smaller avalanches falling down until the base and, thus, flattening the entire sandpile (long range correlation, typical of transitional states). The chain reaction can be imagined as a "branching" process, potentially invading large part of the pile (the size of the avalanche can be easily estimated in terms of number of grains involved); in short, each grain falls down until a position of rest, during the slide the grain hits other grains causing small, and then large, avalanches. It was experimentally demonstrated (Bak, 1991) that, even huge avalanches (posited that we continue to add a grain at a time) keep invariant the slope of the sandpile, given that the probability that a single grain stops is balanced by the probability of a new avalanche. It is worth noting that even huge avalanches invading the entire sandpile happen by a local mechanism (domino effect): each grain only interacts with its neighbour, this is exactly what we observed by RQA as applied to gene disposition along the chromosomes.

If the slope of the sandpile is lower than the critical slope (sub-critical state) the pile will grow until it reaches the critical threshold, if the slope is too steep (super-critical state), the size of the avalanches will be greater than in the critical state so lowering the slope of the pile down to the critical threshold. Thus, the "critical state" attracts both sub-critical and super-critical states. The distribution of the size of avalanches follows a power spectrum distribution: very few large avalanches involving the whole pile, very frequent smaller ones involving only parts of it. In any case, no external observer can predict how, where (in which part of the sandpile) and when a catastrophic avalanche will take place even because the (rare) catastrophic avalanches are 'caused' by added grains as small as those provoking very small events. Such "catastrophic" events depend on the past history of the sandpile and not by the strength of the applied stimulus. This makes SOC completely different by other transitions crucially depending on the driving force (control parameter) and question the usual linear relation between the size of the stimulus and its consequences.

Summarizing, the main properties showed by a system governed by SOC are:
(a) Spatial self-similarity (no characteristic length, small and huge avalanches are both typical of the nature of the 'stable criticality' of the system, i.e. they pertain to the same statistical distribution).
(b) Temporal persistence (memory effects, the "catastrophic" event depends on its history and not on the applied stimulus).
(c) Long-term divergent, correlations (small avalanches together can cause one "catastrophic" event by finding their way across by domino effect).
The domino effect characterizing SOC makes it extremely intriguing for a scientist that wish to provoke a large avalanche, potentially creating a huge restructuring on the system, or to reach a "super-critical" state in which the probability of large avalanches is maximized. The supercritical state (exactly like in domino), is attained by increasing the density of contacts between the elements (domino tablets) so that a small stimulus can trigger a long and branched chain reaction. Nevertheless, we cannot actively drive such a phenomenon but only setting the conditions fostering the appearance of such chain reaction, whose shape and size will depend only by the structure and history of the system at hand.

#### 1.2 References Appendix A


Bak, P., Tang, C., & Wiesenfeld, K. (1987). *Physical review letters*, 59(4), 381.
Bak, P., Tang, C., & Wiesenfeld, K. (1988). *Physical review A*, 38(1), 364.
Bak P. Chen K. (1991) *Scientific American*, 1991: 46-

#### 2 Appendix B: Essential Summary of SOC results on MCF-7 cell response

This Appendix summarizes essential results on self-organized control of genome expression on HRG- and EGF-stimulated MCF-7 breast cancer cells (Nagashima, 2007). HRG stimulation provokes a cell fate transition while EGF does not.

Whole genome expression is dynamically self-organized through the emergence of a critical point (CP). Figure S2.1 shows that overall gene expression is stochastic, however its stochastic dynamics follows that of the center of mass (CM) of the whole expression, called coherent-stochastic behaviors (CSBs) (Tsuchita, 2014). The CSB emerges in *N* ensemble genes with *N*>50 (Tsuchita, 2014): the converging toward a stable barycenter (CM) at increasing gene number is a proof of coherent-stochastic behavior. This convergence reveals that the dynamics of the CM of genome expression describes an attractor of the dynamics of stochastic expression.

##### 2.1 Supplementary Figures Appendix B

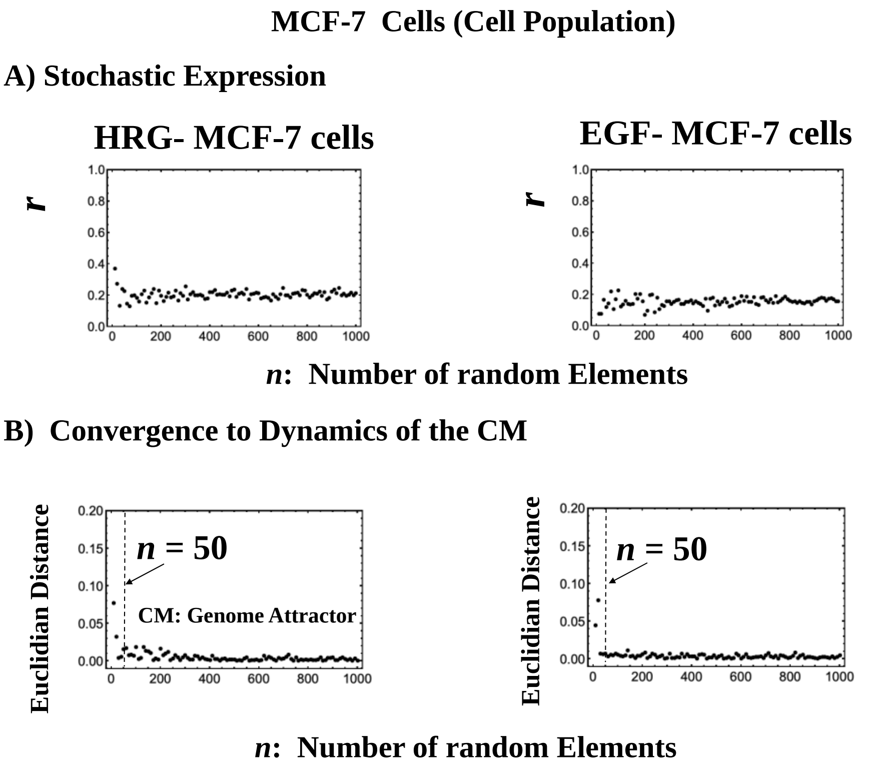


**Figure S2.1: The Center of Mass (CM) of Genome Expression acting as Genome-Attractor.** A bootstrap simulation approach to catch two basic signatures of coherent-stochastic behaviors (CBS): **A)** Stochastic behavior of gene expression shows (relatively low) Pearson correlation (*r*) convergence within randomly selected gene ensembles with 200 repetitions as the number of elements (*n*) is increased. **B)** Convergence of the CM of randomly selected group of expression to the CM of whole expression over experimental time points with 200 repetitions: the dynamics of the CM of randomly selected group (Euclidian distance with the temporal points of the CM of genome expression) converges to that of the CM of the whole expression as the number of selected genes increases (*x*-axis). With A), the CSB emerges in *N* ensemble genes with *N* > 50.

A change in the CP provokes a genome-wide avalanche over the entire genome expression: whole expression follows the change in the CP, the origin of the coherent gene expression behavior spanning the entire genome (Tsuchiya, 2007).

To grasp the mechanism of genome-wide avalanche, Figure S2.2 shows that the CP correspond to the CM of genome expression according to *nrmsf*: the CP represents a specific set of critical genes, which has an activated (ON) or deactivated (OFF) state and furthermore, acts as genome-attractor (Figures S2.2 I-II). The change in critical transition on its singular behaviors (Figures S2.2 III) provides the direct evidence of the ON-OFF state of the CP.

This result implies that in OFF state of the CP, the stochastic perturbations propagate locally, but when the particularity of the disturbance activates the CP, the perturbation can spread over the entire system in a highly cooperative manner (Figure S2.3 VI). As the system approaches its critical point (ON: the CP) where global behavior emerges in a self-organized manner. This is possible in such way that an autonomous critical-control genomic system (genome-engine: Figure S2.3 V) is developed through the formation of dominant cyclic flux between critical states (Figure S2.3 IV). The cell-fate decision is guided by coherent perturbation on the genome engine through the activation of the CP.

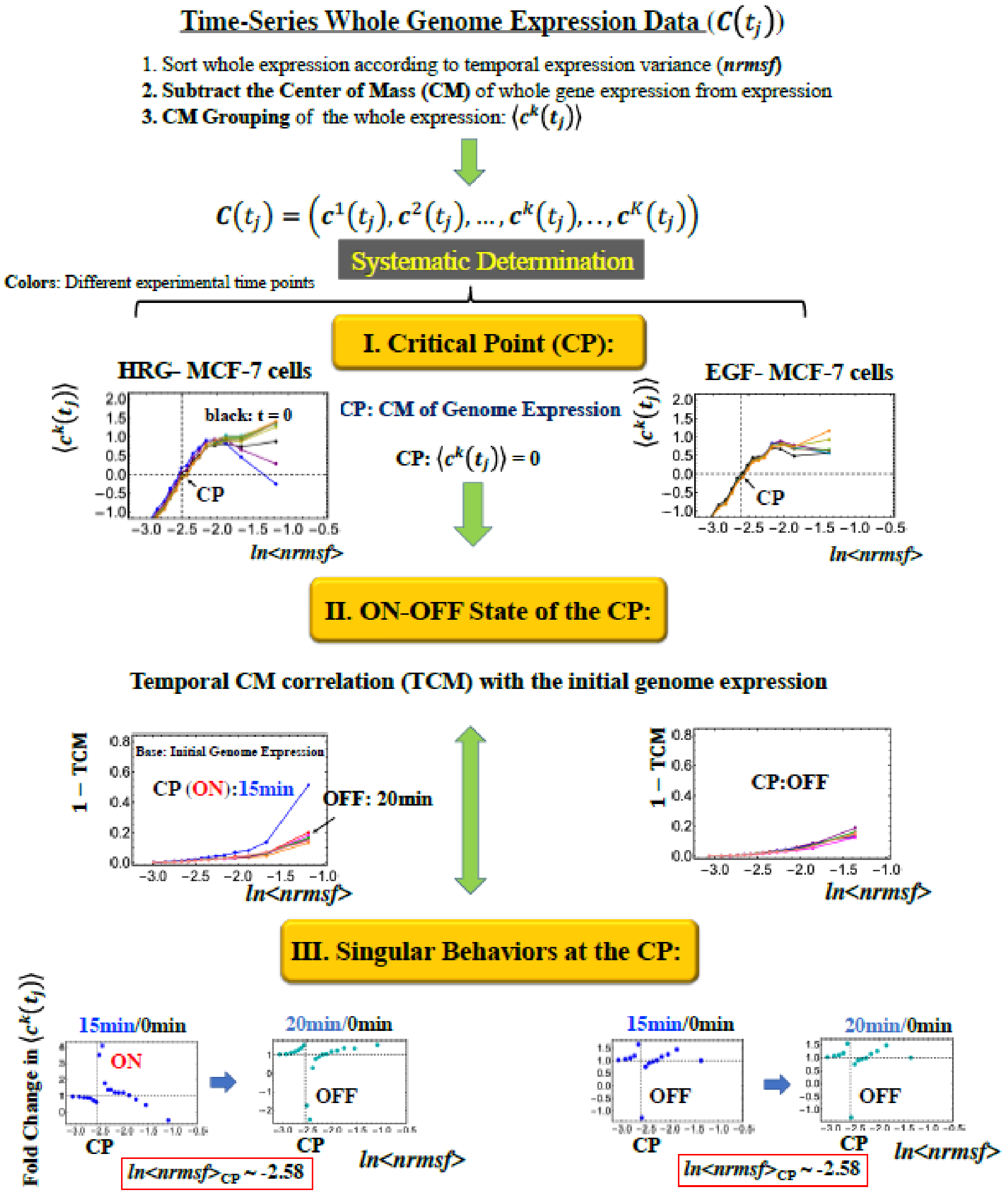


**Figure S2.2: Essential Summary of SOC results of MCF-7 Cells (I-III)**
I) The critical point (CP) corresponds to the CM of genome expression according to the degree of *nrmsf* when each expression is subtracted by the expression value of the CM at a specific time point. Furthermore, the CP correspond to a fixed *nrmsf* value (see more in III) below). The CP stems from a sandpile critical point, where the divergence between up- and down-regulation occurs (refer to Appendix A for sandpile criticality). The CP corresponds to zero-expression point in the CM grouping, which shows that the CP is a specific set of genes corresponding to the CM. Thus, with Figure S2.1, the CP acts as genome-attractor, where a change in the CP provokes a genome-wide avalanche over the entire genome expression: whole expression follows the change in the CP. II) The CP possesses ON-OFF state. This is shown by the grouping (baseline as the CM($t_j$)) according to the degree of *nrmsf*, called ***CM grouping***, $c^k(t_j)$ ($k^{th}$ group; $k$ =1,.., $K$). Temporal development of the CM correlation between the initial and other experimental time point reveals a divergent behavior at $t_j$ = 15min (CP: ON; OFF after 20min) for HRG, whereas for EGF, there are no a divergent behavior over experimental time points (CP: OFF). III) A direct evidence of ON-OFF state in terms of singular behavior at the CP (fold

change in CM grouping) is shown, where for HRG-stimulation, discrete transition of DNA (15min (ON): swelled coil state; 20min (OFF): compact globule state) occurs at the CP, whereas in the case with the EGF-stimulation, such transition does not occur during the early time points.
Note: The occurrence of the singular behavior estimates that average value of $ln(nrmsf)$ of the CP indicates $ln(nrmsf)_{CP}$ = -2.58 for both HRG and EGF.

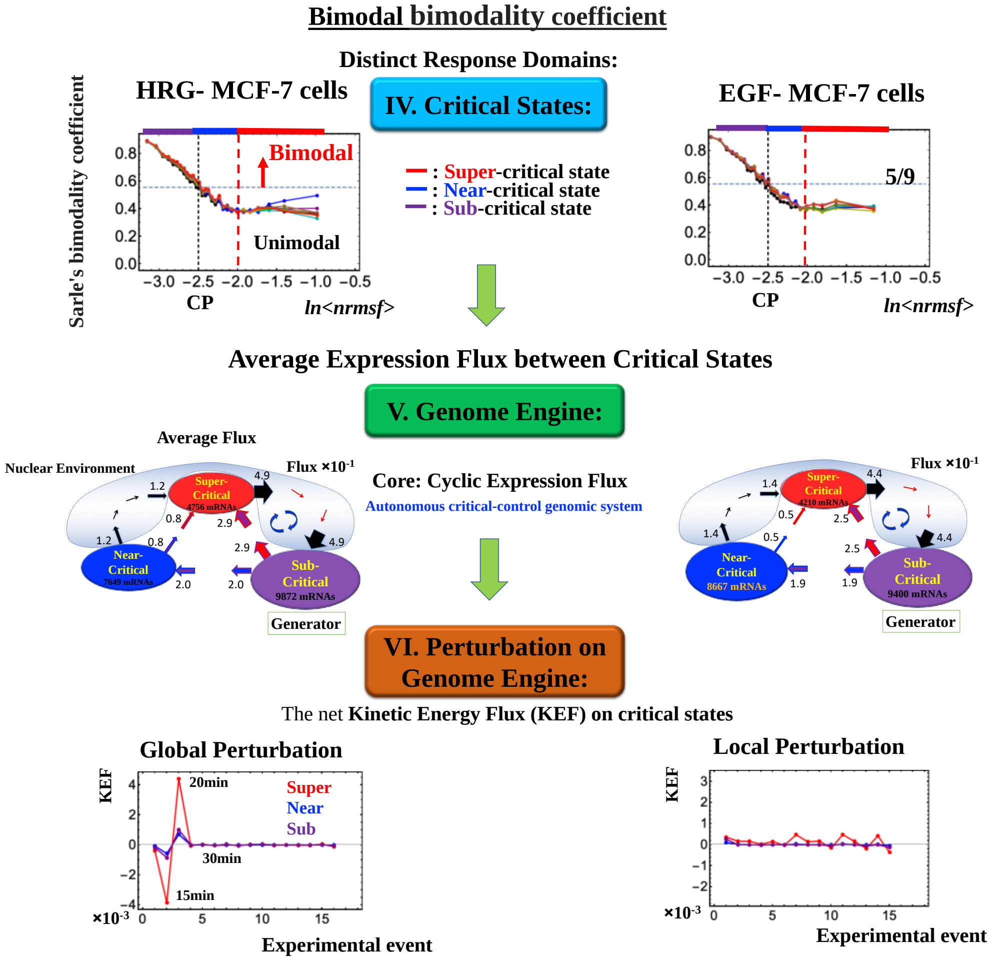


**Figure S2.3: Essential Summary of SOC results of MCF-7 Cells (IV-VI)**
IV) The Sarle's bimodal coefficient ($b$) of CM grouping shows that at the CP, bimodal transition ($b$> 5/9) occurs, and three distinct domains (critical states): unimodal- transit -bimodal transition. V)Genome-engine is shown by average expression flux between critical states through cell nuclear environment: Sub-Super cyclic flux forms a dominant flux flow to establish the genome engine mechanism. The sub-critical state acts as a "large piston" for short moves (low-variance expression) and the super-critical state as a "small piston" for large moves (high-variance expression). The "ignition switch" corresponds to a critical point (the genome-attractor) and the 'pistons' are connected through a dominant cyclic state flux as a "camshaft", resulting in the anti-phase dynamics of two pistons (see theoretical foundation for expression flux analysis in (Tsuchiya, 2016, 2018, 2019). VI)The net kinetic-energy-flux (KEF) reveals that HRG-stimulation induces global perturbation on the genome-engine (involvement of critical states) for HRG occurs at 15-20min (exactly coinciding with the timing of ON-OFF of the CP), whereas EGF-stimulation induces only vivid activation in super-critical, i.e., local perturbation, where the stochastic perturbations propagate locally. Coherent perturbation on the genome-engine through the activation of the CP guides the cell-fate decision.

#### 2.2 References Appendix B

Nagashima, T., Shimodaira, H., Ide, K., Nakakuki, T., Tani, Y., Takahashi, K., Yumoto, N., Hatakeyama, M. (2007) Quantitative transcriptional control of ErbB receptor signaling undergoes graded to biphasic response for cell differentiation. *J Biol Chem* 282: 4045–4056.

Tsuchiya, M., Hashimoto, M., Takenaka, Y., Motoike, I. N., Yoshikawa, K. (2014). Global genetic response in a cancer cell: Self-organized coherent expression dynamics. *PLOS One* 9: e97411.

Tsuchiya, M., Giuliani, A., Hashimoto, M., Erenpreisa, J., & Yoshikawa, K. (2015). Emergent Self-Organized Criticality in gene expression dynamics: Temporal development of global phase transition revealed in a cancer cell line. *PLOS One*, 10(6), e0128565.

Tsuchiya, M., Wong, ST., Yeo, ZX., Colosimo, A., Palumbo, MC., Crescenzi, M., Mazzola, A., Negri, R., Bianchi, MM., Selvarajoo, K., Tomita, M., Giuliani, A. (2007). Gene expression waves: cell cycle independent collective dynamics in cultured cells. *FEBS J*. 274, 2874-2886.

Tsuchiya, M., Giuliani, A., Hashimoto, M., Erenpreisa, J., Yoshikawa, K. (2016). Self-organizing global gene expression regulated through criticality: Mechanism of the cell-fate change. PLOS ONE 11: e0167912;

Tsuchiya, M., Giuliani, A., Yoshikawa, K. (2018). A Quantitative Evaluation of Symmetry Breaking In Nonlinear-Oscillatory System - Based on Flux Dynamics (Effective force) View Point. Presentation; doi: 10.13140/RG.2.2.34048.74240;

Tsuchiya, M., Giuliani, A., Yoshikawa, K. (2019). Underlying Genomic Mechanism for Cell-Fate Change from Embryo to Cancer Development Preprint, bioRxiv: doi: https://doi.org/10.1101/637033

#### 3 Appendix C: Recurrence quantification analysis (RQA)

Recurrence quantification analysis (RQA) is a nonlinear technique, originally developed by Eckmann et al. (Eckmann, 1987) as a purely graphical method and then made quantitative by Webber and Zbilut (1994). It was successfully applied to different fields ranging from physiology (Zimatore, 2000) molecular dynamics (Manetti, 1999) and economics (Orlando, 2018). The concept of recurrence is straightforward: for any ordered series (time or spatial), a recurrence is simply a point which repeats itself. This makes RQA immediately applicable to biopolymers like DNA (Pizzi, 2001) and proteins (Giuliani, 2002) where the order of monomers (nucleotides, amino-acid residues, genes as in our case) along the sequence is isomorphic to a time series sampled at discrete equally spaced intervals. Recurrences are the most basic of relations shaping a given system, since they are strictly local and independent of any mathematical assumption regarding the system itself. Furthermore, it is worth stressing that calculation of recurrences, unlike other methods such as Fourier, Wigner-Ville, or wavelets, requires no transformation of the data and does not require a minimum length of the series.
RQA works on the embedding matrix (EM) of an original linear time series. EN is a n-dimensional matrix generated by shifting the original series of a fixed lag. In the case of the discrete series:
10, 11, 21,32, 41, 35, 40, 19..., the corresponding 4-dimensional EM becomes:
10 11 21 32
11 21 32 41
21 32 41 35
32 41 35 40
41 35 40 19
35 40 19
40 19
19

The rows of the embedding matrix (EM) correspond to subsequent windows of length 4 (embedding dimension) along the sequence. RQA is based on the computation of the Euclidean distance matrix (DM) between the rows (epochs) of the EM, looking for epochs close to each other (recurrences).
The concept of a recurrence can be expressed as follows: given a reference point, X0, and a ball (Br) of radius r, a point X is said to recur (with reference to X0) if

$$Br(X0) = \{X: | X - X0|\} \leq r$$

In the case of a time series, i.e., of a system occupying in different times different positions along a trajectory in a suitable state space, the recurrences correspond to the time points where the system passes nearby to already visited states. In our case, in which the elements are single gene expressions, time corresponds to the location of genes along the chromosome and the recurrences are patches, with a length equal to the embedding dimension, sharing their expression profile with other patches along the chromosome. The number and relative positions of recurrences are expressed by recurrence plots (RP) that are symmetrical N* N arrays in which a point is placed at (i, j) whenever a point Xi on the trajectory is close to another point Xj. The closeness between Xi and Xj is expressed by calculating the Euclidian distance between these two normed vectors, i.e., by subtracting one from the other. The two points are scored as recurrent if their Euclidian distance is lower than the pre-defined fixed radius r. This procedure allows for the construction of the Recurrence Plot (RP) correspondent to the distance matrix between the different epochs (rows

of the embedding matrix) filtered, by the action of the radius, to a binary 0/1 matrix having a 1 (dot) for distances falling below the radius and a 0 for distances greater than radius.
Because graphical representations may be difficult to evaluate, Zbilut and Webber (Webber, 1994) developed several strategies to quantify features of such plots. Hence, the quantification of RPs leads to the generation of some global descriptors as:

REC (percent of plot filled with recurrent points).
DET (percent of recurrent points forming diagonal lines with a minimum of 2 adjacent points).
ENT (Shannon information entropy of the line length distribution).
MAXLINE, length of longest line segment (the reciprocal of which is an approximation of the largest positive Lyapunov exponent and is a measure of system divergence.
TREND (measure of the paling of recurrent points away from the central diagonal).
These recurrence descriptors quantify the deterministic structure and complexity of the plot.

In our case, the original series were the subsequent standardized gene expression values along the chromosome at each time point. Each chromosome was separately analysed obtaining practically identical results for all the chromosomes at any time point. The chosen embedding dimension was set to 3 and the radius was set to 40% of mean distance.

#### 3.1 Supplementary Figure Appendix C

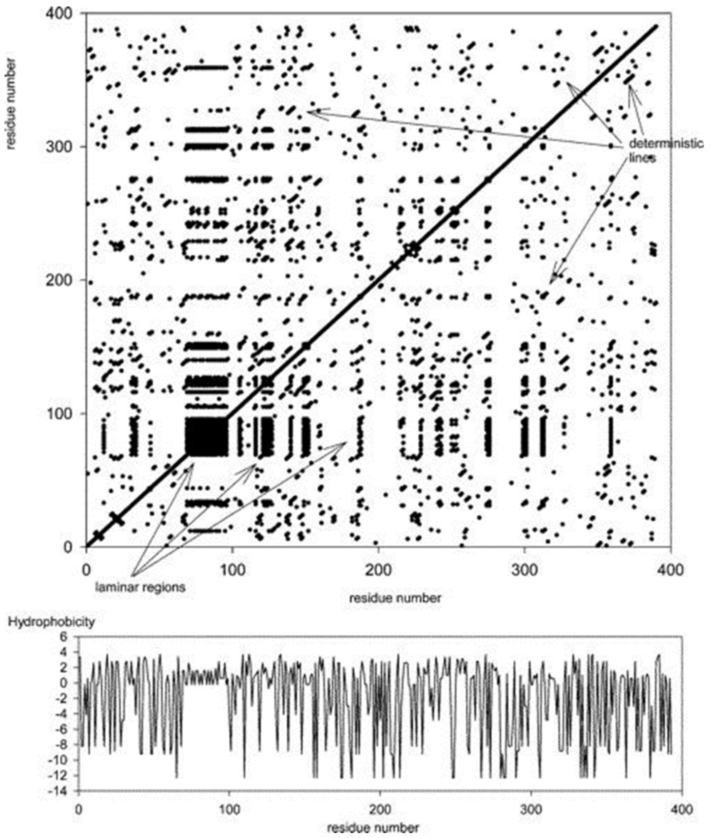


**Figure S3.1**

Figure S3.1 reports an exemplar recurrence plot (hydrophobicity patterning along P53 protein sequence, together with the original series that generated it).

#### 3.2 References Appendix C


Eckmann, J. P.; Kamphorst, S. O.; Ruelle, D. (1987) *Europhys. Lett.* 4, 324- 358.

Manetti, C.; Ceruso, M. A.; Giuliani, A.; Webber, C. L.; Zbilut, J. P. (1999), *Phys. Rev. E* 59, 992.

Giuliani, A., Benigni, R., Zbilut, J. P., Webber, C. L., Sirabella, P., Colosimo, A. (2002). *Chemical Reviews*, 102(5), 1471-1492.

Orlando, G., & Zimatore, G. (2018) *Chaos, Solitons & Fractals*, 110, 82-94.

Pizzi, E., & Frontali, C. (2001). *Genome Research*, 11(2), 218-229.

Webber, C. L.; Zbilut J. P (1994). *J. Appl. Physiol*., 76, 965.

Zimatore, G., Giuliani, A., Parlapiano, C., Grisanti, G., & Colosimo, A. (2000), *Journal of Applied Physiology*, 88(4), 1431-1437.
